# Autophagy-dependent filopodial kinetics restrict synaptic partner choice during *Drosophila* brain wiring

**DOI:** 10.1101/762179

**Authors:** Ferdi Ridvan Kiral, Gerit Arne Linneweber, Svilen Veselinov Georgiev, Bassem A. Hassan, Max von Kleist, Peter Robin Hiesinger

## Abstract

Brain wiring is remarkably precise, yet most neurons readily form synapses with incorrect partners when given the opportunity. Dynamic axon-dendritic positioning can restrict synaptogenic encounters, but the spatiotemporal interaction kinetics and their regulation remain essentially unknown inside developing brains. Here we show that the kinetics of axonal filopodia restrict synapse formation and partner choice for neurons that are not otherwise prevented from making incorrect synapses. Using 4D imaging in developing *Drosophila* brains, we show that filopodial kinetics are regulated by autophagy, a prevalent degradation mechanism whose role in brain development remains poorly understood. With surprising specificity, autophagosomes form in synaptogenic filopodia, followed by filopodial collapse. Altered autophagic degradation of synaptic building material quantitatively regulates synapse formation as shown by computational modeling and genetic experiments. Increased filopodial stability enables incorrect synaptic partnerships. Hence, filopodial autophagy restricts inappropriate partner choice through a process of kinetic exclusion that critically contributes to wiring specificity.

## Introduction

Synapse formation and synaptic partner choice are based on molecular and cellular interactions of neurons in all animals ^1–5^. Brain wiring diagrams are highly reproducible, yet most, if not all, neurons have the ability to form synapses with incorrect partners, including themselves ^6,7^. During neural circuit development, spatiotemporal patterning restricts when and where neurons ‘see each other’ ^8–10^. Positional effects can thereby prevent incorrect partnerships, even when neurons are not otherwise prevented from forming synapses ^7,11,12^. When and where neurons interact with each other to form synapses is a fundamentally dynamic process. Yet, the roles of neuronal interaction dynamics, e.g. the speed or stability of filopodial interactions, is almost completely unknown for dense brain regions in any organism. Our limited understanding of the dynamics of synaptogenic encounters reflects the difficulty to observe, live and *in vivo*, synapse formation at the level of filopodial dynamics in intact, normally developing brains ^13,14^.

Fly photoreceptors (R cells) are the primary retinal output neurons that relay visual information with highly stereotypic synaptic connections in dense brain regions, namely the lamina and medulla neuropils of the optic lobe ^15–17^. Intact fly brains can develop in culture, enabling live imaging at the high spatiotemporal resolution necessary to measure photoreceptor axon filopodial dynamics and synapse formation throughout the entire developmental period of circuit assembly ^13,14,18^. Axonal filopodia inside the developing brain stabilize to form synapses through the accumulation of synaptic building material, but how limiting amounts of building material in filopodia are regulated is unknown ^14^.

Macroautophagy (autophagy hereafter) is a ubiquitous endomembrane degradation mechanism implicated in neuronal maintenance and function ^19^. Neuronal autophagy has been linked to neurodegeneration ^20^ as well as synaptic function in the mature nervous system ^21,22^. Comparably little is known about developmental autophagy in the brain. Functional neurons develop in the absence of autophagy ^19,23,24^. In specific neurons in worms and flies, loss of autophagy leads to reduced synapse development ^25,26^. By contrast, in the mouse brain loss of autophagy in neurons leads to increased dendritic spine density due to defective pruning after synapse formation ^27,28^. Despite numerous links to neurodevelopmental disorders, it remains unknown if and how developmental autophagy can contribute to synaptic partner choice and circuit connectivity, especially in dense brain regions.

In this study, we show that loss of autophagy in *Drosophila* photoreceptor neurons leads to increased synapse formation and the recruitment of incorrect postsynaptic partners. Autophagy directly and selectively regulates the kinetics of synaptogenic axon filopodia, a phenotype that could only be revealed through live observation during intact brain development. Autophagic modulation of the kinetics of synaptogenic filopodia restricts what neurons ‘see each other’ to form synapses, thereby critically contributing to the developmental program that ensures synaptic specificity during brain development.

## Results

We have previously observed the formation of autophagosomes at the axon terminals of developing photoreceptor neurons R1-R6 in the developing *Drosophila* brain, but their function has remained unknown ^29^. Previous analyses of loss of autophagy in fly photoreceptors have not revealed any obvious developmental defects ^24,30,31^.

### Flies with autophagy-deficient photoreceptors exhibit increased neurotransmission and visual attention behavior

To probe for previously undetected synaptic defects, we blocked autophagy in developing photoreceptor neurons using molecularly well-defined mutants for the essential autophagy proteins Atg7 and Atg6 (fly homolog of Beclin-1) ^24,30^. We validated loss of the key autophagosome marker Atg8 in both *atg7* and *atg6* mutants (Supplementary Fig. 1a-b’ and d). Rescue of *atg6* with the photoreceptor-specific driver GMR-Gal4 reversed this effect and led to a significant increase in Atg8-positive compartments compared to wild type (Supplementary Fig. 1c-c’ and d).

As expected, the eyes and axonal projections of photoreceptor neurons mutant for *atg6* or *atg7* in otherwise wild type brains exhibited no obvious defects in fixed preparations (Fig. 1a, b). Photoreceptor neurons are known to exhibit neurodegeneration with ageing ^31^. To assay photoreceptor function directly following autophagy-deficient development, we therefore recorded electroretinograms (ERGs) from the eyes of newly eclosed flies. Autophagy-deficient photoreceptors exhibited normal depolarizing responses to light, indicating functional phototransduction and healthy neurons (Fig. 1c, d). Surprisingly, ‘on’ transient amplitudes, which are indicative of synaptic transmission and the ability to elicit a postsynaptic response, were increased 30-50% in both mutants (Fig. 1c, e). Conversely, increased autophagy in transgenically rescued *atg6* photoreceptors reversed this effect and resulted in a significant reduction of ‘on’ transients (Fig. 1c, e).

**Fig. 1.**
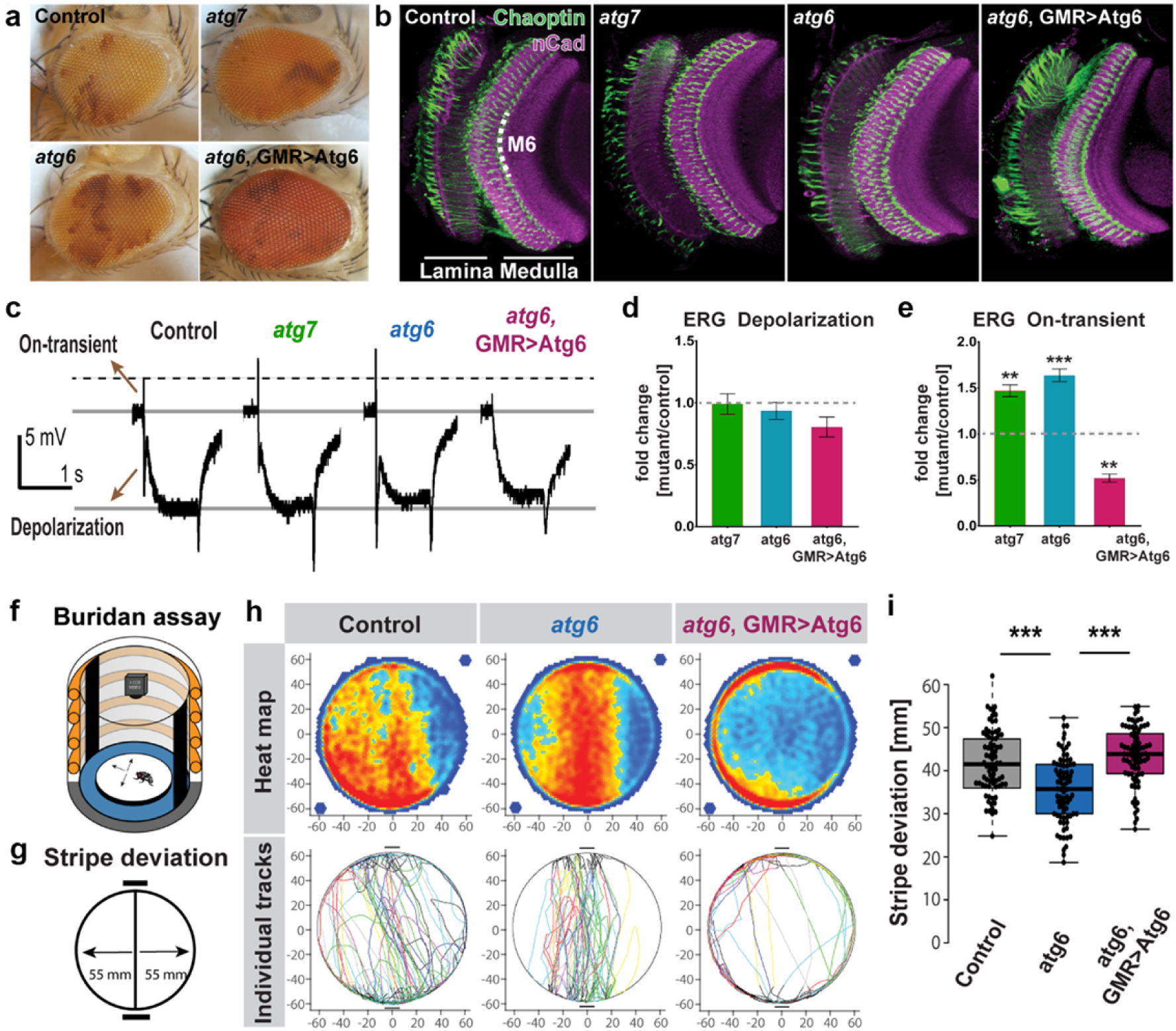
Autophagy deficiency in *Drosophila* photoreceptors leads to increased neurotransmission and visual attention. **a-b,** Newly-hatched (0-day old) genetic mosaic flies with autophagy-deficient (*atg6* and *atg7* mutants) photoreceptors exhibit normal eye morphology (**a**) and axonal projections in the optic lobe (**b**). **c,** Representative electroretinogram (ERG) traces. **d-e,** Quantification of ERG depolarization (**d**) and on-transient (**e**) amplitudes relative to control. Rescue of *atg6* mutant photoreceptors with GMR>atg6 expression leads to overcompensation and increased autophagy (see Supplementary Fig. 1). n=20 flies per condition. Unpaired t-test; *p<0.05, **p<0.01, ***p<0.001. Error bars denote mean ± SEM. **f,** Buridan’s paradigm arena to measure object orientation response of adult flies, with two black stripes positioned opposite to each other as visual cues. **g,** The parameter ‘stripe deviation’ measures how much a fly deviates from a straight path between the black stripes in the arena. **h,** Stripe fixation behavior of adult flies with *atg6* mutant photoreceptors, photoreceptors with upregulated autophagy (*atg6*, GMR>Atg6) and their genetically matched controls are shown on the population level (heatmap) and as individual tracks. Flies with *atg6* mutant photoreceptors show reduced stripe deviation, whereas increased autophagy (*atg6*, GMR>Atg6) leads to increased stripe deviation. **i,** Quantification of stripe deviation. n=60 flies per condition, two-way ANOVA and Tukey HSD as post hoc test, ***p<0.001.

Next, we asked whether loss of autophagy selectively in photoreceptors affected fly vision. We used the simple visual choice assay Buridan’s paradigm, in which wing-clipped flies walk freely in a circular, uniformly illuminated arena with two high contrast black stripes placed opposite to each other (Fig. 1f) ^32^. In this assay, flies with functional vision walk back and forth between the two high contrast objects. We chose the parameter ‘stripe deviation’, which measures how much a single fly deviates from an imaginary line between two black stripes, as a behavioral read-out of visual attention (Fig. 1g). Flies with *atg6* or *atg7*-deficient photoreceptors were assayed and compared to their genetic background matched controls. Surprisingly, in both mutants the flies with autophagy-deficient photoreceptors exhibited increased visual attention behavior (decreased stripe deviation) compared to their genetically matched controls (Fig. 1h, i and Supplementary Fig. 2). Increased autophagy in *atg6* rescued photoreceptors reversed this effect again in an overcompensatory manner similar to ERG responses (Fig. 1h, i). We conclude that flies with photoreceptors that developed in the absence of autophagy can see, but their vision is characterized by both increased neurotransmission and increased visual attention.

### Autophagy-deficient Drosophila photoreceptors form supernumerary synapses

To assess whether the alterations in neurotransmission and vision were due to altered numbers of synapses, we generated sparse clones of photoreceptors R1-R6, and R7 expressing the active zone marker GFP-Brp^short^. This marker specifically localizes to presynaptic active zones without affecting synaptic development or function and is suitable for live imaging ^14,33^. Loss of autophagy resulted in a 25%-80% increase in synapse numbers, while increased autophagy in rescued *atg6* mutant photoreceptors reversed this effect and significantly reduced synapse numbers (Fig. 2a-f). Photoreceptors R1-R6 form columnar terminals in a single layer neuropil, whereas R7 axon terminals span six morphologically distinct layers and form the majority of synapses in the most proximal layer M6 ^17,34^. We were therefore surprised to see many supernumerary synapses in autophagy-deficient R7 axon terminals at more distal layers M1-M3 (Fig. 2g; red boxes in Fig. 2a-d’). These putative synapses along the distal shaft of autophagy-deficient R7 axons were stable based on live imaging of Brp^short^- labelled active zones with 15 min resolution over several hours at P70 (70% pupal development; Supplementary Movie 1). Brp stability is indicative of mature synapses and suggests that ectopic Brp puncta in fixed images are not the consequence of axonal transport defects or defective synaptic capture of Brp-positive transport vesicles. These observations raised the question whether loss of autophagy leads to genuine supernumerary synapses and, if so, whether these would be formed with correct postsynaptic partners.

**Fig. 2.**
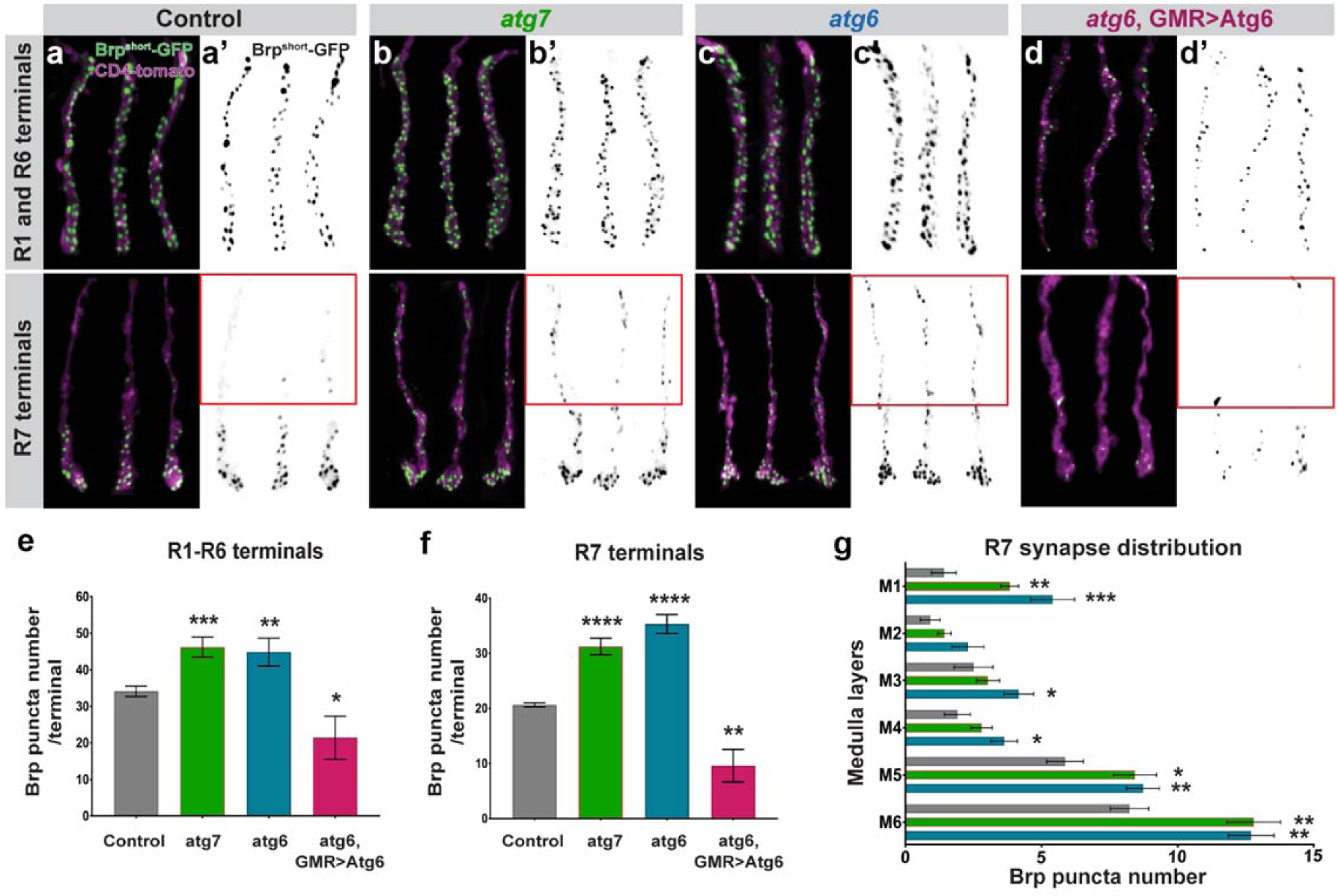
Autophagy-deficient *Drosophila* photoreceptors form supernumerary synapses. **a-d’,** Representative images of R1-R6 and R7 photoreceptor axon terminals with Brp^short-^GFP marked active zones in wild-type **(a-a’)**, *atg7* mutant **(b-b’)**, *atg6* mutant **(c-c’)**, and *atg6*, GMR>Atg6 **(d-d’)**. Red boxes show supernumerary synapses in loss of autophagy at distal part of R7 axon terminals. **e-f,** Number of Brp puncta per terminal in R1-R6 (**e**) and R7 (**f**) photoreceptors. n=40 terminals per condition. Unpaired t-test; *p<0.05, **p<0.01, ***p<0.001, ****p<0.001. Error bars denote mean ± SEM. **g,** Number of Brp puncta in distinct medulla layers along R7 axon terminals (See ‘Materials and Methods’ for the definition of medulla layers). Unpaired t-test; *p<0.05, **p<0.01, ***p<0.001. Error bars denote mean ± SEM.

### Autophagy-deficient R7 photoreceptors contact incorrect postsynaptic neurons

The synaptic partners of R7 photoreceptors have been quantitatively characterized based on EM reconstruction of several medulla columns, revealing highly stereotypic connections ^17^. The main post-synaptic target of R7 photoreceptors is the wide-field amacrine neuron Dm8 ^17,35^. Apart from Dm8s, R7s form fewer connections with Tm5 neuron subtypes that have dendritic fields spanning from M3 to M6 ^34,35^. To identify the postsynaptic partners of autophagy-deficient R7 photoreceptors, we used the recently developed anterograde trans-synaptic tracing method ‘*trans*-Tango’, which labels post-synaptic neurons for a given neuron without a need for previous knowledge about the nature of the connections ^36^. We used an R7-specific driver (Rhodopsin4-Gal4) and restricted its expression to mutant R7 photoreceptors, while all other neurons, including all postsynaptic partners, are wild type. Consistent with known post-synaptic targets of R7s, *trans*-Tango with wild-type R7s mainly labelled Dm8s and Tm5s (Fig. 3a). By contrast, loss of autophagy in R7s led to more widespread labelling of post-synaptic neurons (Fig. 3b) and an overall increase of the number of postsynaptically connected cells, as expected for supernumerary functional synapses (Fig. 3c). Through application of a sparse-labeling protocol of *trans*-Tango, we further identified several cell types, including Mi1, Mi4, Mi8, Tm1, C2, and C3 that are not normally postsynaptic to R7 based on connectome data ^15,17,37,38^ (Fig. 3d, e). Mi1 and Mi4, for example, are part of the motion-detection pathway, to which R7 is not known to provide input ^39,40^. Notably, the number of individual neurons detected for these six ectopically connected neurons correlated distinctly with the position of their presumptive dendritic trees: Mi1, C3 and C2 were most often labeled and all three have presumptive dendrites in layers M1 and M5 (Fig. 3e, f) ^41^; most ectopic R7 synapses were detected in layer M1, M5 and M6 (Fig. 2g); at the other end of the spectrum, Mi8 and Tm1 were both 4-5fold less often detected and have presumptive dendrites in layer M2 and M3, where we counted fewer ectopic synapses (Fig. 2g and Fig. 3e, f) ^41^. These findings suggest that the postsynaptic neurons labeled by *trans*-Tango are incorrect partners connected through axon-dendritic contacts with R7.

**Fig. 3.**
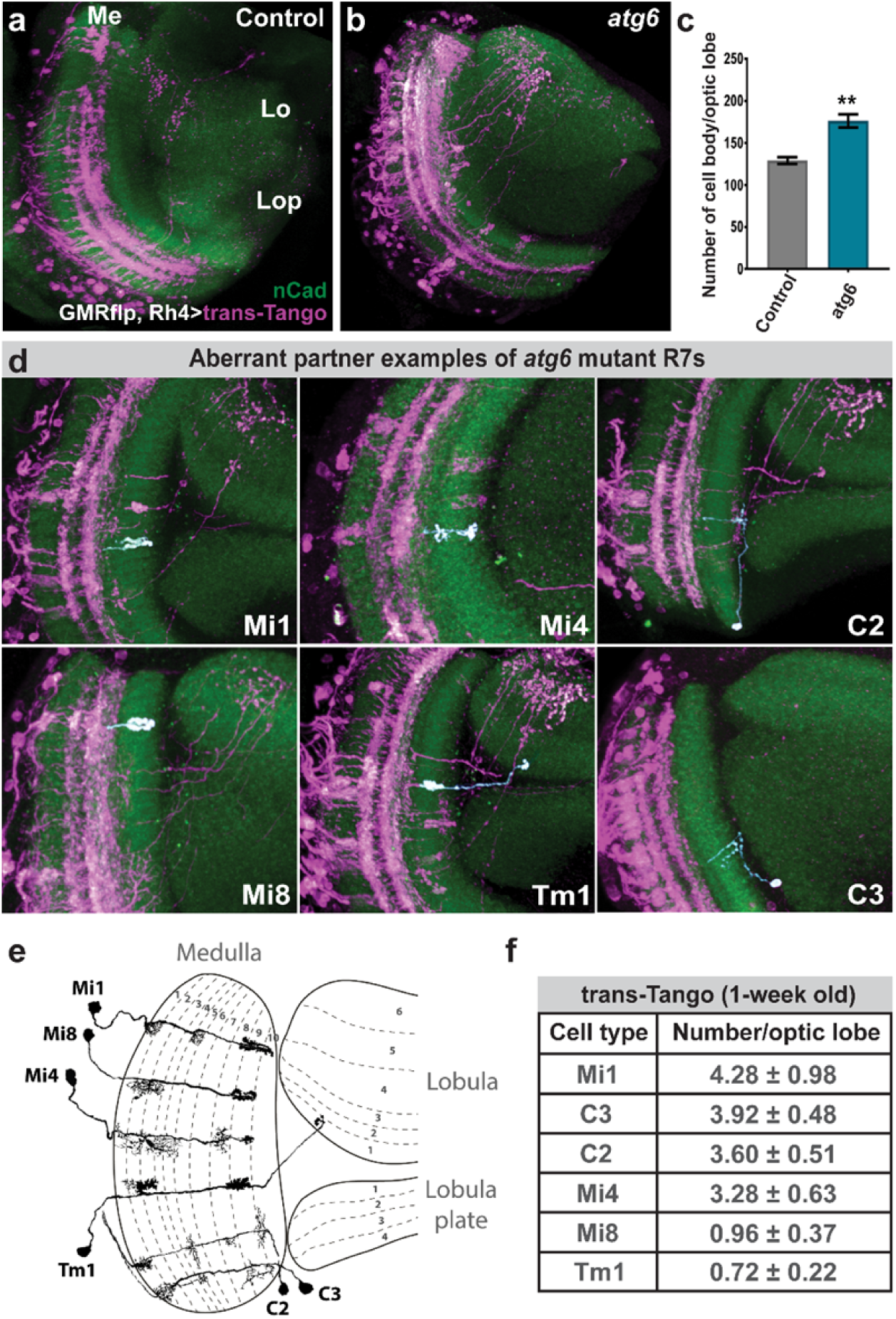
Loss of autophagy leads to synaptic connections with aberrant neuronal partners. **a-b,** Neurons post-synaptic to control (**a**) and *atg6* mutant (**b**) R7s are labelled with *trans*-Tango (see ‘Materials and Methods’ for full genotypes, magenta=post-synaptic neurons, green=CadN; Me=medulla, Lo=lobula, Lop=Lobula plate). **c,** Number of post-synaptic neurons per optic lobe for control and *atg6* mutant R7s based on trans-Tango-labeled cell body counts. Unpaired t-test; **p<0.01. **d,** Examples of aberrant neuronal partners of autophagy-deficient R7s, with individual neurons pseudo-colored in white. **e**, Schematic of dendritic and axonal arborization of aberrant neuronal partners (Adapted from Fiscbach and Dittrich, 1989)^41^. **f**, Number of each aberrant neuronal partners per optic lobe from 1-week old fly brains. Note that only *∼*10% of R7s are mutant for *atg6* and trans-Tango labeling is dependent on synaptic strength between partners and progressively increase through age. See ‘Materials and Methods’ for detailed *Drosophila* genotypes used to perform trans-Tango experiments.

### Synapses with incorrect postsynaptic neurons are functional based on activity-dependent GRASP

To test whether these contacts are functional synapses, we next used the activity-dependent GRASP method (GFP reconstitution across synaptic partners), which is based on trans-synaptic complementation of split GFP only when synaptic vesicle release occurs ^42,43^. Based on available cell-specific driver lines and the underlying genetics, we could test three of the ectopic pairs identified with *trans*-Tango: potential synapses between R7 and Mi1, C2 or Mi4. For all three cases, wild type neurons rarely showed isolated synaptic signals (Fig. 4a-c’). In contrast, *atg6* mutant photoreceptors formed abundant synapses in all three cases (Fig. 4d-f’). These findings also indicate that the *trans*-Tango results were not due to an effect of altered autophagy on the ectopically expressed proteins of the trans-Tango system. We conclude that loss of autophagy in R7 photoreceptor terminals leads to ectopic synapse formation with inappropriate postsynaptic neurons.

**Fig. 4.**
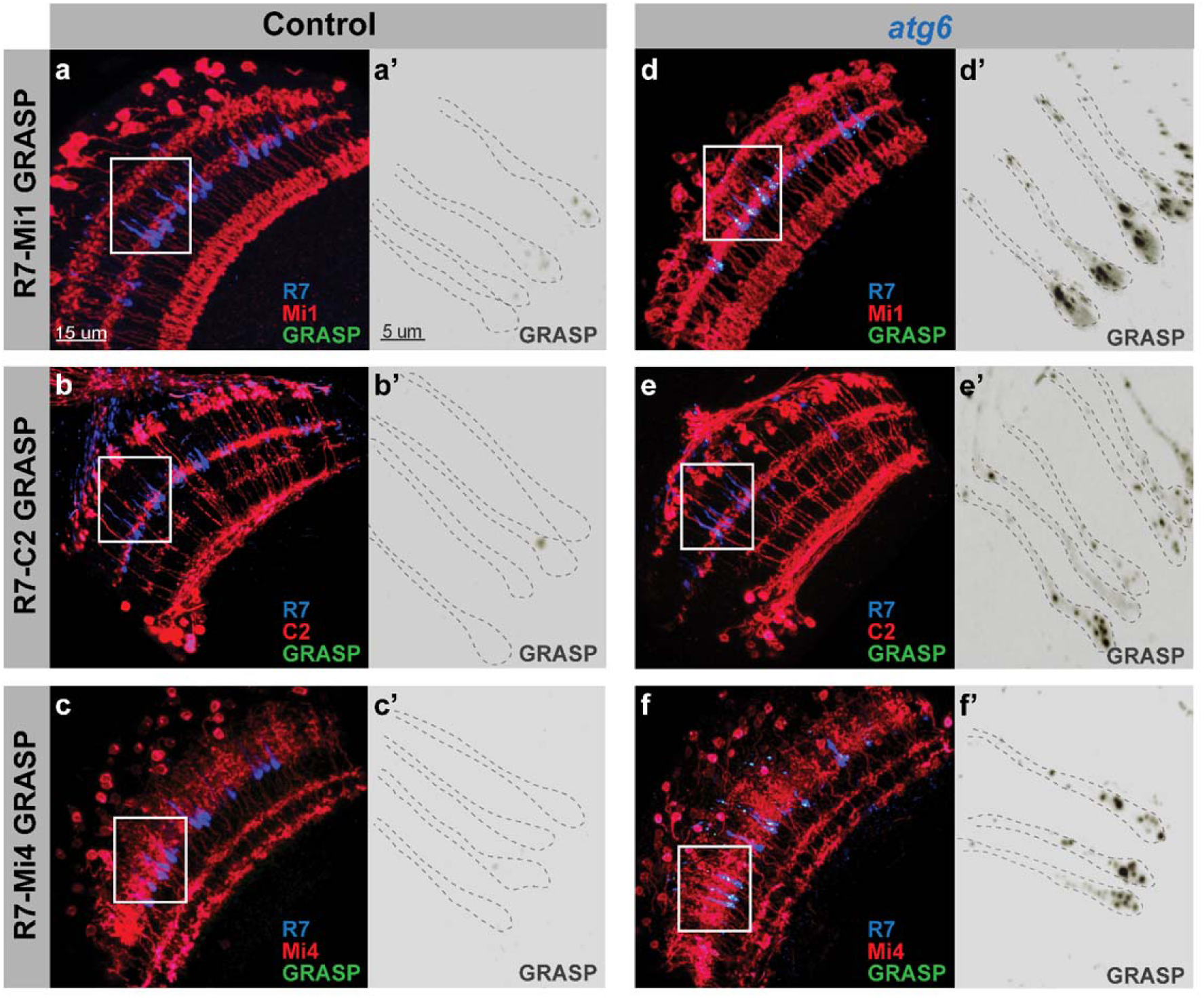
Synaptic connections between autophagy-deficient R7s and aberrant postsynaptic partners are functional based on activity-dependent GRASP. **a-c’,** Activity-dependent GRASP between control R7s and Mi1s (**a-a’**), C2s (**b-b’**), and Mi4s (**c-c’**) show that wild-type R7s very rarely form synaptic connections, if any, with Mi1, C2, and Mi4 neurons. **d-f’,** Activity-dependent GRASP between *atg6* mutant R7s and Mi1s (**d-d’**), C2s (**e-e’**), and Mi4s (**f-f’**) show widespread active synaptic connections between autophagy-deficient R7s and aberrant post-synaptic partners. Regions inside yellow rectangles are shown in close-up images as single greyscale GRASP channels. See ‘Materials and Methods’ for Mi1, Mi4, and C2-specific LexA drivers and detailed *Drosophila* genotypes used to perform GRASP experiments.

Taken together, we conclude that loss of autophagy in photoreceptors does not affect overall axon terminal morphology and transmission of visual input, but selectively leads to increased synapse formation, which includes inappropriate postsynaptic partners, and increased visual attention behavior. But how does defective autophagy at the developing pre-synapse affect synaptic partner choice mechanistically?

### Autophagy modulates the stability of synaptogenic filopodia

To test when and where exactly autophagosomes function during synapse formation, we performed live imaging experiments of autophagosome formation in developing R7 axon terminals in developing brains. Autophagosomes have previously been shown to form at axon terminals in vertebrate primary neuronal cell culture using the temporal series of autophagosome maturation reporters Atg5-GFP (early) and Atg8-GFP (late) ^44^. Surprisingly, we found autophagosome formation based on these probes selectively at the rare, bulbous tips of synaptogenic filopodia of R7 axon terminals, followed by filopodial collapse (Fig. 5a; Supplementary Fig. 3; Supplementary Movie 2).

**Fig. 5.**
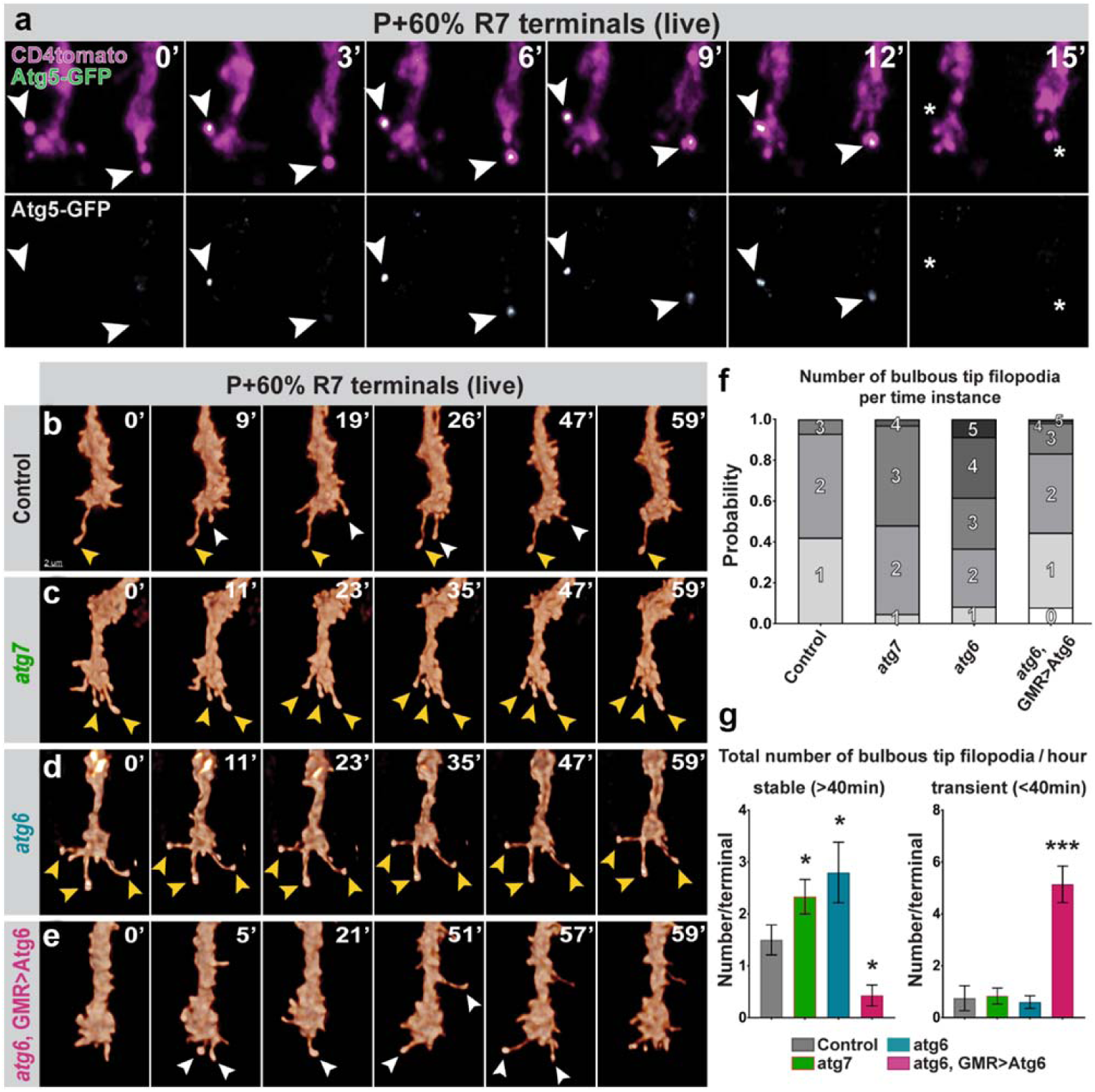
Autophagy regulates the stability of synaptogenic filopodia at axon terminals. **a,** Live imaging of Atg5-GFP expressing R7 axon terminals in intact, developing *Drosophila* brain shows formation of autophagosomes at the bulbous tips of synaptogenic filopodia ^14^ followed by the collapse of filopodia (P+60%). **b-e,** Live imaging of R7 axon terminals at P+60% (during synaptogenesis) revealed increased stability of synaptogenic filopodia in autophagy-deficient R7 terminals (**c** and **d**) and decreased stability in R7 terminals with upregulated autophagy (**e**) compared to control (**b**). Yellow arrowheads: stable synaptogenic filopodia; white arrowheads: unstable bulbous tip filopodia. **f,** Number of concurrently existing bulbous tip filopodia per R7 axon terminal per time instance. **g,** Total number of synaptogenic filopodia per R7 axon terminal per hour. Autophagy-deficient R7 terminals exhibit significantly more stable synaptogenic filopodia (>40min) whereas upregulated autophagy leads to filopodia destabilization. n=7 terminals per condition. Unpaired t-test; *p<0.05, ***p<0.001. Error bars denote mean ± SEM.

We have recently shown that altered numbers of synaptogenic filopodia lead to changes in synapse numbers ^14^. We therefore tested the effects of a loss of autophagy on R7 axon terminal filopodial dynamics during synapse formation (developmental time point P60). Both *atg6* and *atg7* mutants exhibited selectively increased lifetimes of the population of long-lived axonal filopodia compared to wild type and *atg6* rescued photoreceptors (Supplementary Fig. 4; Supplementary Table 1). Wild type axon terminals only formed 1-2 synaptogenic filopodia, as characterized by their bulbous tips, at any point in time (Fig. 5b, f-g), which previously led us to propose a serial synapse formation process that slowly spreads out the formation of 20-25 synapses over 50 hours ^14^ (also see Supplementary Movie 3). In contrast, loss of *atg6* or *atg7* in R7 axon terminals led to 3-4 synaptogenic filopodia at any time point (Fig. 5c-d and 5f-g; Supplementary Movie 3). As expected for synaptogenic filopodia, almost all supernumerary bulbous tips were stable for more than 40 minutes (Fig. 5g). Increased autophagy in *atg6* rescued mutant photoreceptors reversed this effect and lead to a significant reduction and destabilization of synaptogenic filopodia (Fig. 5e-g; Supplementary Movie 3). Consistent with selective autophagosome formation in synaptogenic filopodia tips, the changes to filopodial dynamics were remarkably specific to long-lived, synaptogenic filopodia (Fig. 5b-g; Supplementary Fig. 4; Supplementary Table 1). In sum, analyses of R7 axon terminal dynamics during synapse formation in the intact brain revealed autophagosome formation in synaptogenic filopodia and a specific effect of autophagy function on the kinetics and stability of these filopodia.

### A developmental model quantitatively predicts the measured increase in synapse numbers based on measured filopodial kinetics in autophagy mutants

Next we asked whether the observed changes to the kinetics of synaptogenic filopodia are sufficient to quantitatively explain changes in synapse formation throughout the second half of fly brain development. We first counted the numbers of overall filopodia, bulbous tip filopodia and synapses at time points every ten hours between P40 and P100 in fixed preparations (Fig. 6a-c). Compared to control, loss of *atg6* or *atg7* in photoreceptors led to mild increases in overall filopodia, while leaving the rates of change largely unaltered between time points (Fig. 6a). In contrast, numbers of synaptogenic bulbous tip filopodia are increased 2-fold throughout the main period of synapse formation (P60-P80; Fig. 6b; Supplementary Fig. 5). Synapse numbers, based on presynaptic *Brp^short^* labeling, commences indistinguishably from wild type, but then increases at a higher rate throughout brain development (Fig. 6c).

**Fig. 6.**
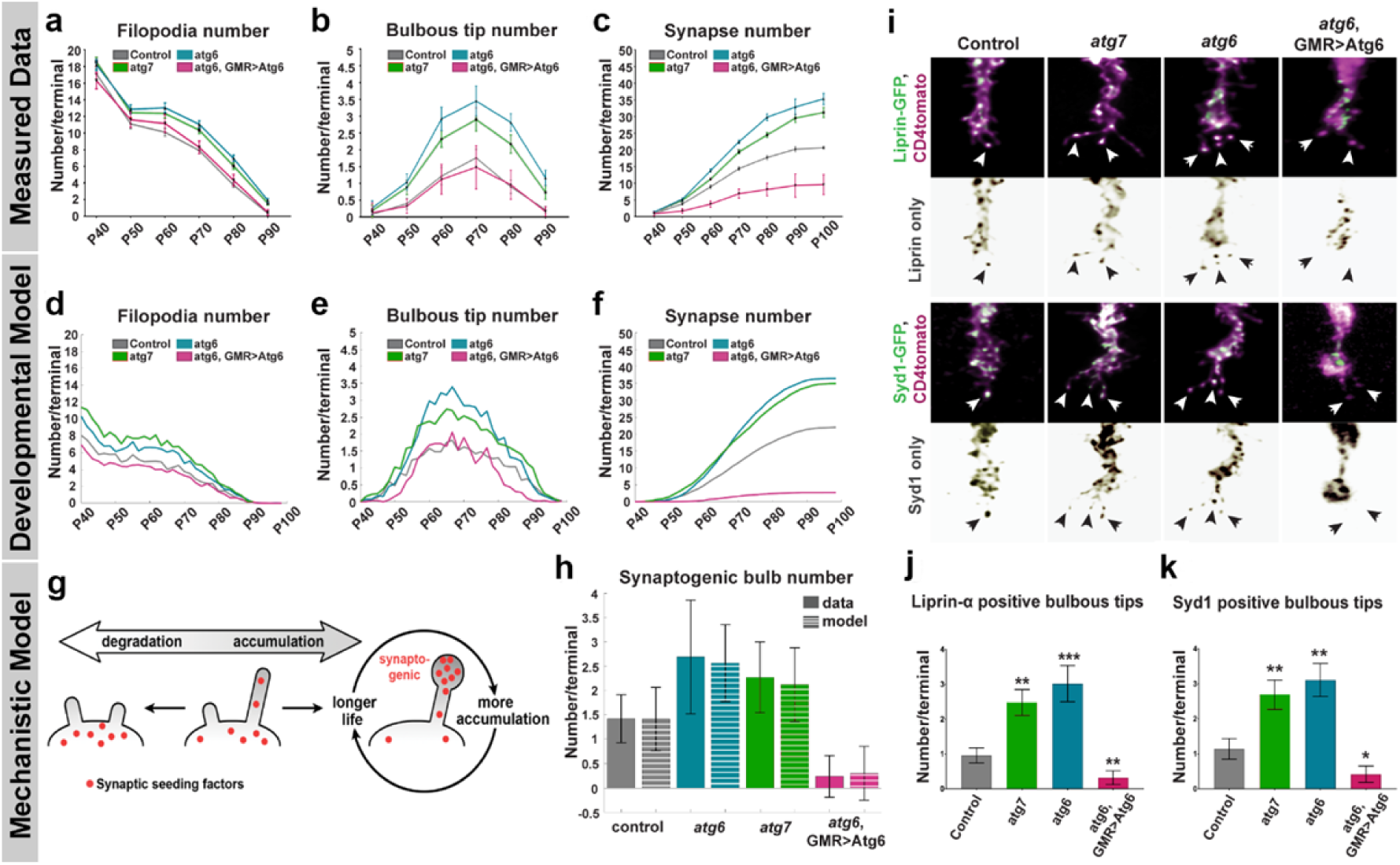
Loss of autophagy increases the number of synaptogenic filopodia through defective synaptic seeding factor degradation, leading to increased synapse formation throughout development. **a-c,** Quantification of filopodia numbers (**a**), synaptogenic filopodia numbers (**b**), and Brp puncta numbers (**c**) during synaptogenesis (P40-P90) per R7 axon terminal based on fixed data. n=40 terminals per condition. **d-f,** Markov State Model simulation based on data in (**a**) and live data at P+60% (Figure 5) for filopodia numbers (**d**), synaptogenic filopodia numbers (**e**), and Brp puncta numbers per R7 axon terminal (**f**). **g,** The mechanistic model: accumulation of synaptic seeding factors stabilizes synaptogenic filopodia; autophagic degradation of synaptic seeding factors destabilizes filopodia. **h,** Measured (solid bars) and simulated (striped bars) synaptogenic filopodia numbers at P+60% (the simulated data are based on synaptic seeding factor availability, see Supplementary Fig. 6). **i,** Representative images of synaptic seeding factors (Syd1 and Liprin-α) localizing to synaptogenic filopodia. **j-k,** Quantifications of the number of Liprin-α (**j**) and Syd1 (**k**) positive synaptogenic filopodia. n=30 terminals per condition. Unpaired t-test; *p<0.05, **p<0.01, ***p<0.001. Error bars denote mean ± SEM.

We previously developed a data-driven Markov state model that predicts the slow, serial development of synapses throughout the second half of brain development based on stochastic filopodial exploration and one-by-one selection of synaptogenic filopodia ^14^. To test how autophagy-dependent changes of filopodial kinetics affect synapse formation in the model, we used the measured live dynamics of filopodia at P60 (Fig. 5b-g; Supplementary Fig. 4; Supplementary Tables 1-3) together with the measured fixed time points data for filopodia (Fig. 6a-b; Supplementary Fig. 5) as input. As shown in Fig. 6d to 6f, the model recapitulates all aspects of synaptogenic filopodial dynamics and synapse formation for both loss and upregulation of autophagy. The model thereby shows that the measured changes in filopodial kinetics, and specifically altered stabilization of synaptogenic filopodia, are sufficient to cause the observed alterations in synapse formation over time (see ‘Mathematical modeling’ in Materials and Methods). These findings raise the question how autophagy can specifically regulate the kinetics of synaptogenic filopodia mechanistically.

### Degradation of synaptic building material through autophagy modulates filopodia kinetics and synapse formation

We have previously shown that the early synaptic seeding factors Syd1 and Liprin-α are allocated to only 1-2 filopodia at any given time point and that their loss leads to the destabilization of synaptogenic filopodia and a loss of synapses ^14^. Autophagy is a protein degradation pathway that affects filopodia stability in opposite ways in loss-versus gain-of-function experiments. We therefore hypothesized that autophagic degradation may directly regulate the availability of synaptic building material in filopodia. We first tested this idea using a second Markov state model that simulates the stabilization of filopodia as a function of seeding factor accumulation and degradation on short time scales (Fig. 6g and Supplementary Fig. 6a). In this ‘winner-takes-all’ model, synaptic seeding factors are a limiting resources in filopodia that increase filopodia lifetime, which in turn increases the time available for further accumulation of synaptic seeding factors, creating a positive feedback loop ^14^. If autophagy plays a role in the degradation of synaptic seeding factors, then decreased autophagic degradation of synaptic seeding factors should lead to more synaptogenic filopodia, while increased autophagic degradation should reduce synaptogenic filopodia through further restriction of the limiting resource (Fig. 6g and Supplementary Fig. 6a). The simulations show that the measured number of synaptogenic filopodia (Fig. 6h) and their lifetimes (Supplementary Fig. 6) can be quantitatively explained by degradation, and thus availability, of synaptic seeding factors for both loss and upregulation of autophagy at P60. Specifically, the number of long-lived filopodia at autophagy-deficient axon terminals was increased compared to control and conversely increased autophagic activity led to a decreased lifespan of filopodia as measured (Supplementary Fig. 4 and Supplementary Table 1). Hence, the mechanistic model predicts that modulation of autophagy affects the degradation and availability of synaptic seeding factors. This primary defect causes secondary changes to filopodial kinetics and synapse formation.

To validate the primary defect, we expressed GFP-tagged versions of the synaptic seeding factors Syd-1 and Liprin-α and analyzed their restricted localization to synaptogenic filopodia. Autophagy-deficient terminals contain 2-3 times more synaptogenic filopodia with synaptic seeding factors compared to control; conversely, upregulation of autophagy leads to reduction of seeding factors in filopodia (Fig. 6i-k). In addition, the majority of Atg8-positive autophagosomes present at filopodia tips colocalizes with with Syd-1 and Liprin-α (Supplementary Fig. 7a-c). Previous work in primary vertebrate neuronal culture as well as *Drosophila* R1-R6 photoreceptors has shown that autophagosomes formed at axon terminals traffic retrogradely to the cell body ^29,44^. We therefore analyzed photoreceptor cell bodies and found large Atg8-positive multivesicular bodies containing Syd-1 (Supplementary Fig. 7d-e). Together, these findings indicate that autophagy controls the amount of synaptic seeding factors in filopodia, thereby affecting their stability and potential to form synapses.

### Incorrect synaptic partnerships result from terminal-wide loss of kinetic restriction, not from filopodia-specific regulation of autophagy

Autophagy-dependent filopodial kinetics and synapse formation could lead to synapses with incorrect partners through at least two mechanisms. In one scenario, autophagy could be triggered only in specific filopodia, e.g. based on a molecular signal for a contact with an incorrect partner neuron in a wrong layer. Loss of autophagy would then lead to a defect in the specific removal of incorrect synapses. In support of this idea, specific presynaptic proteins have recently been shown to induce autophagy at specific places in the presynapse ^45,46^. Alternatively, autophagy could set a global threshold for kinetic restriction, such that only synaptic partners with sufficient spatial availability and molecular affinity can form synapses.

To distinguish between these two models, we quantified the relative increases of all filopodia, synaptogenic filopodia, and synapses along the R7 axon terminal in medulla layers M1-M6 (Fig. 7a-d). Loss of either *atg6* or *atg7* increases the absolute numbers of synaptogenic filopodia and synapses in all medulla layers equally approximately 1.5-fold (dotted lines in Fig. 7b-d). As a result, the relative levels of synaptogenic filopodia and synapses between layers M1-M6 remain the same as in wild type (solid lines in Fig. 7b-d). These data indicate that autophagy is not differentially triggered in filopodia in specific medulla layers. Instead, loss of autophagy equally increases the stability of synaptogenic filopodia across the R7 terminal, resulting in the stabilization of only few filopodia in layers with low baseline filopodial activity, and more pronounced increases in layers with higher baseline filopodial activity. Conversely, destabilization of filopodia along the entire R7 axon terminal in wild type effectively excludes synapse formation in layers with few filopodia, e.g. in layer M2 (Fig. 7a-d). We conclude that autophagy levels set a threshold for kinetic restriction across the R7 axon terminal.

**Fig. 7.**
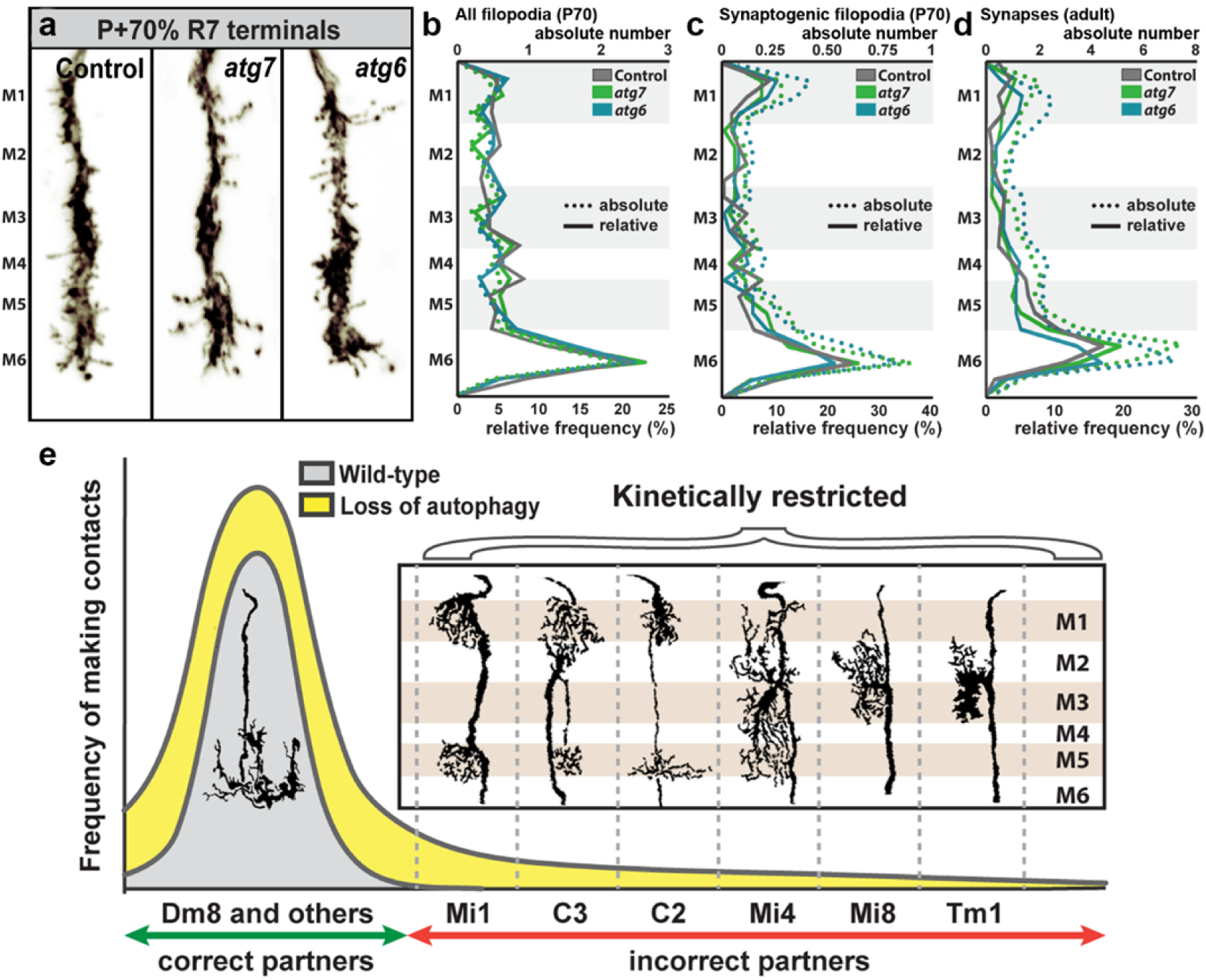
Loss of autophagy recruits incorrect synaptic partners by lowering an axon terminal-wide threshold for kinetic restriction of synapse formation. **a,** Representative R7 axon terminals at P+70% with medulla layer information. Note that the edge of medulla (M0) is defined as 0 and the end of M6 layer is defined as 100 to calculate relative positions of all filopodia and bulbous tip filopodia and distributed to medulla layers (M1-M6) using the relative thickness of medulla layers defined by Fiscbach and Dittrich, 1989^41^. **b-d,** Relative frequency (solid lines) and absolute numbers (dotted lines) of all filopodia at P+70% (**b**), synaptogenic filopodia at P+70% (**c**), and synapses at 0-day old adult (**d**). M1-M6 denote medulla layers. n=40 terminals per condition. **e,** Model: Loss of autophagy during synaptogenesis increases the probability distribution (yellow area) compared to wild type (grey area) of forming connections with post-synaptic partners through increased filopodial stability. Note that cells with projections at medulla layers where R7s form most of their synapses (Mi1, Mi4, C2, C3) incorrectly synapse with R7s with higher probability than the cells with projections at medulla layers where R7s form a few, if any, synapses (Mi8, Tm1) (See Figure 3e and 3f).

The threshold for kinetic restriction effectively excludes synapse formation with at least six potential postsynaptic partners that are not otherwise prevented from forming synapses with R7 (Fig. 7e). We note that the localization of the presumptive dendritic trees of these six neuron types correlates well with the probabilities to be incorrectly recruited as postsynaptic partners (Fig. 3f and Fig. 7e). We speculate that specificity arises through a combination of context-dependent molecular interactions, positional effects and kinetic restriction.

## Discussion

Brain wiring requires synaptic partner choices that are both specific and robust in time and space ^47^. To what extent spatiotemporal vicinity of potential partner neurons facilitates or determines partner choice remains unclear. Our findings suggest that spatiotemporal vicinity is restricted by filopodial kinetics and that axon terminal autophagy functions as a modulator of these dynamics. Hence, kinetic restriction of synaptogenic filopodia is a means to effectively exclude synapse formation with incorrect partners (Fig. 7e). Conversely, increased stabilization of synaptogenic filopodia reveals a surprisingly varied population of interneurons that have the principle capacity to form synapses with R7 axon terminals. At least Mi1, Mi4, C3, C2, Mi8 and Tm1 neurons in medulla columns are not prevented by ‘molecular mismatch’ from forming synaptic contacts with R7 in a normal developmental environment.

### Kinetic restriction sharpens synaptic specificity based on molecular and cellular interactions

Numerous studies have shown that neurons in ectopic locations readily form synapses with incorrect partners, including themselves ^6,7,48^. On the other hand, Mi1, Mi4, C3, C2, Mi8 and Tm1 are likely to express cell surface proteins that bias the likelihood of synaptic contacts with R7 and other partners ^10,16,49^. Axonal and dendritic interaction dynamics may greatly facilitate, or restrict, what partner neurons get ‘to see each other’ and initiate synapse formation based on molecular interactions ^1^. Recent evidence highlighted the importance of positional strategies for synaptic partner choice prior to such molecular recognition ^7,11,48^. Here we have shown that positional effects do not only depend on when and where neuronal processes can be seen in fixed preparations, but are a function of their dynamics and stabilization kinetics. Hence, synaptic specificity can emerge from the context-dependent combination of molecular interactions with a cell biological mechanism like autophagy, that by itself carries no synaptic specificity information. We speculate that different nenal thresholds for kinetic restriction can critically contribute to sharpen specificity as part of the brain’s developmental growth program.

### A role for developmental autophagy in synapse formation and brain wiring

Our data support the idea that autophagy indiscriminately destabilizes R7 synaptogenic filopodia in a manner consistent with the local degradation of a limiting resource of proteins required for synapse formation. Specificity of autophagic degradation can be triggered through interactions with proteins that themselves serve as cargo or restrict the time and place where potentially less specific engulfment occurs ^19,45,46^. The bulbous tips of synaptogenic filopodia are a small space that may be easily destabilized through autophagic engulfment of proteins and other cargo, even if that engulfment were to occur in a non-selective manner. We therefore propose that putative cargo-specificity of autophagy may not be a prerequisite for the developmental function of autophagy described here.

We have previously shown that spatiotemporally regulated membrane receptor degradation is required for synapse-specific wiring in the *Drosophila* visual system ^50^. In order for degradation of receptors or synaptic seeding factors to serve as regulators of spatiotemporal specificity, the degraded proteins must undergo continuous turnover. Specificity therefore arises through a combination of developmentally regulated protein synthesis, trafficking and degradation, which are likely to differ for different proteins and neurons at different points in time and space.

Based on this combinatorial model for specificity, we speculate that many mutations and single nucleotide polymorphisms in the genome can result in small cell biological changes that differentially affect neurons during brain wiring. The changes effected through such modulatory, ‘permissive’ mechanisms may not be predictable at the level of circuit wiring and behavior, yet they can cause meaningful changes to behavior that are both selectable and heritable and thus a means of evolutionary programming of neural circuits.

## Acknowledgements

We would like to thank Mathias Wernet and all members of the Hiesinger, Wernet and Hassan labs for their support and helpful discussions. We thank Eric Baehrecke and Stephan Sigrist for reagents. F.R.K acknowledges Gizem Sancer for her help to design ‘*trans*-Tango’ experiments. This work was supported by the NIH (RO1EY018884) and the German Research Foundation (DFG, SFB 958, SFB186) and FU Berlin. B.H. was supported by an Einstein BIH Fellowship. M.v.K acknowledges financial support from the German ministry for education and science (BMBF) through grant number 031A307 and from the Einstein Stiftung Berlin and the DFG, provided through the excellence cluster Math+.

## Author contributions

F.R.K and P.R.H. designed the project. F.R.K performed all experiments except the Buridan’s paradigm. Behavioral experiments were designed, carried out and analyzed by G.A.L and B.A.H. S.V.G. performed 4D tracking analyses of filopodial dynamics. M.v.K. performed all computational modeling. F.R.K, B.A.H., M.v.K and P.R.H. wrote the paper.

## Competing interests

The authors declare no competing interests.

## MATERIALS AND METHODS

### EXPERIMENTAL MODEL AND SUBJECT DETAILS

Flies were reared at 25°C on standard cornmeal/yeast diet unless stated otherwise. For developmental analyses white pre-pupae (P+0%) were collected and incubated at 25°C to pupal stages stated on figures. The following Drosophila strains were either obtained from Bloomington Drosophila Stock Center (BDSC) or other groups: *atg6*^1^ and UAS-Atg6.ORF.3xHA (E.H. Baehrecke); *atg7*^d4^ (T.Neufeld); UAS-Brpshort-GFP, UAS-Syd1-GFP, and UAS-Liprinα-GFP (S.Sigrist); Trans-tango flies (G.Barnea); GRASP flies (BDSC); ey3.5flp, GMRflp, GMR-Gal4, FRT42D, FRT82B, GMR-Gal80, tub-Gal80, UAS-CD4-tdGFP, UAS-CD4-tdtomato, UAS-Atg5-GFP, UAS-Atg8-GFP, GMR22F08-LexA (C2-specific driver), GMR49B06-LexA (Mi4-specific driver), and GMR19F01-LexA (Mi1-specific driver) (BDSC).

#### Drosophila genotypes

##### Figure 1

**a-i**, <u>Controls</u>: ey3.5flp; FRT42D/FRT42D, Cl^w+^, ey3.5flp; GMR-Gal4/+; FRT82B/FRT82B, Cl^w+^, *<u>atg7</u>*: ey3.5flp; FRT42D, *atg7*^d4^/FRT42D, Cl^w+^, *<u>atg6</u>*: ey3.5flp;GMR-Gal4/+; FRT82B, *atg6*^1^/FRT82B, Cl^w+^, <u>*atg6*, GMR>Atg6</u>: ey3.5flp;GMR-Gal4/+; FRT82B, *atg6*^1^, UAS-Atg6.ORF.3xHA /FRT82B, Cl^w+^.

##### Figure 2

**a-g**, <u>Controls</u>: GMRflp; FRT42D, GMR-Gal80/FRT42D; GMR-Gal4, UAS-CD4-tdtomato/UAS-Brp^short^-GFP, GMRflp; GMR-Gal4, UAS-CD4-tdtomato/UAS-Brp^short^-GFP; FRT82B/FRT82B, tub-Gal80, *<u>atg7</u>*: GMRflp; FRT42D, GMR-Gal80/FRT42D, *atg7*^d4^; GMR-Gal4, UAS-CD4-tdtomato/UAS-Brp^short^-GFP, *<u>atg6</u>*: GMRflp; GMR-Gal4, UAS-CD4-tdtomato/UAS-Brp^short^-GFP; FRT82B, *atg6*^1^/FRT82B, tub-Gal80, *<u>atg6</u>*<u>, GMR>Atg6</u>: GMRflp; GMR-Gal4, UAS-CD4-tdtomato/UAS-Brp^short^-GFP; FRT82B, *atg6*^1^, UAS-Atg6.ORF.3xHA/FRT82B, tub-Gal80.

##### Figure 3

**a-f**, <u>Control</u>: GMRflp/UAS-myrGFP, QUAS-mtdtomato(3xHA); Rh4-Gal4/trans-Tango; FRT82B/FRT82B, tub-Gal80, *<u>atg6</u>*: GMRflp/UAS-myrGFP, QUAS-mtdtomato(3xHA); Rh4-Gal4/trans-Tango; FRT82B, *atg6*^1^/FRT82B, tub-Gal80.

##### Figure 4

**a-c’**, <u>Control</u>: GMRflp; Rh4-Gal4, UAS-nSyb::splitGFP1-10, LexAop-splitGFP11::GFP/ GMR19F01-LexA (Mi1) or GMR22F08-LexA (C2) or GMR49B06-LexA (Mi4); FRT82B/FRT82B, tub-Gal80

**d-f’**, *<u>atg6</u>*: GMRflp; Rh4-Gal4, UAS-nSyb::splitGFP1-10, LexAop-splitGFP11::GFP/ GMR19F01-LexA (Mi1) or GMR22F08-LexA (C2) or GMR49B06-LexA (Mi4); FRT82B, *atg6*^1^/FRT82B, tub-Gal80

##### Figure 5

**a**, GMRflp; FRT42D, GMR-Gal80/FRT42D; GMR-Gal4, UAS-CD4-tdtomato/UAS-Atg5-GFP.

**b-g**, <u>Controls</u>: GMRflp; FRT42D, GMR-Gal80/FRT42D; GMR-Gal4, UAS-CD4-tdGFP, GMRflp; GMR-Gal4, UAS-CD4-tdGFP; FRT82B, tub-Gal80/FRT82B, *<u>atg7</u>*: GMRflp; FRT42D, *atg7*^d4^/FRT42D, tub-Gal80; GMR-Gal4, UAS-CD4-tdGFP, *<u>atg6</u>*: GMRflp; GMR-Gal4, UAS-CD4-tdGFP; FRT82B, *atg6*^1^/FRT82B, tub-Gal80, *<u>atg6</u>*<u>, GMR>Atg6</u>: GMRflp; GMR-Gal4, UAS-CD4-tdGFP; FRT82B, *atg6*^1^, UAS-Atg6.ORF.3xHA/FRT82B, tub-Gal80.

##### Figure 6

**a-b**, <u>Controls</u>: GMRflp; FRT42D, GMR-Gal80/FRT42D; GMR-Gal4, UAS-CD4-tdGFP, GMRflp; GMR-Gal4, UAS-CD4-tdGFP; FRT82B, tub-Gal80/FRT82B, *<u>atg7</u>*: GMRflp; FRT42D, *atg7*^d4^/FRT42D, GMR-Gal80; GMR-Gal4, UAS-CD4-tdGFP, *<u>atg6</u>*: GMRflp; GMR-Gal4, UAS-CD4-tdGFP; FRT82B, *atg6*^1^/FRT82B, tub-Gal80, *<u>atg6</u>*<u>, GMR>Atg6</u>: GMRflp; GMR-Gal4, UAS-CD4-tdGFP; FRT82B, *atg6*^1^, UAS-Atg6.ORF.3xHA/FRT82B, tub-Gal80.

**c**, <u>Control</u>: GMRflp; FRT42D/FRT42, GMR-Gal80; GMR-Gal4, UAS-CD4-tdtomato, UAS-Brp^short^-GFP, *<u>atg7</u>*: GMRflp; FRT42D, *atg7*^d4^/FRT42, GMR-Gal80; GMR-Gal4, UAS-CD4-tdtomato, UAS-Brp^short^-GFP, *<u>atg6</u>*: GMRflp; GMR-Gal4, UAS-CD4-tdtomato, UAS-Brp^short^-GFP; FRT82B, *atg6*^1^/FRT82B, tub-Gal80, *<u>atg6</u>*<u>, GMR>Atg6</u>: GMRflp; GMR-Gal4, UAS-CD4-tdtomato/UAS-Brp^short^-GFP; FRT82B, *atg6*^1^, UAS-Atg6.ORF.3xHA/FRT82B, tub-Gal80.

**i-k**, <u>Controls</u>: GMRflp; FRT42D, UAS-Liprin-α-GFP or UAS-Syd-1-GFP/FRT42D, GMR-Gal80; GMR-Gal4, UAS-CD4-tdtomato, GMRflp; GMR-Gal4, UAS-CD4-tdtomato, UAS-Liprin-α-GFP or UAS-Syd-1-GFP; FRT82B/FRT82B, tub-Gal80, *<u>atg7</u>*: GMRflp; FRT42D, *atg7*^d4^, UAS-Liprin-α-GFP or UAS-Syd-1-GFP/FRT42D, tub-Gal80; GMR-Gal4, UAS-CD4-tdtomato, *<u>atg6</u>*: GMRflp; GMR-Gal4, UAS-CD4-tdtomato, UAS-Liprin-α-GFP or UAS-Syd-1-GFP; FRT82B, *atg6*^1^/FRT82B, tub-Gal80; *<u>atg6</u>*<u>, GMR>Atg6</u>: GMRflp; GMR-Gal4, UAS-CD4-tdtomato/ UAS-Liprin-α-GFP or UAS-Syd-1-GFP; FRT82B, *atg6*^1^, UAS-Atg6.ORF.3xHA/FRT82B, tub-Gal80.

##### Figure 7

**a-c**, <u>Controls</u>: GMRflp; FRT42D, GMR-Gal80/FRT42D; GMR-Gal4, UAS-CD4-tdGFP, GMRflp; GMR-Gal4, UAS-CD4-tdGFP; FRT82B, tub-Gal80/FRT82B, *<u>atg7</u>*: GMRflp; FRT42D, *atg7*^d4^/FRT42D, GMR-Gal80; GMR-Gal4, UAS-CD4-tdGFP, *<u>atg6</u>*: GMRflp; GMR-Gal4, UAS-CD4-tdGFP; FRT82B, *atg6*^1^/FRT82B, tub-Gal80.

**d**, <u>Control</u>: GMRflp; FRT42D/FRT42, GMR-Gal80; GMR-Gal4, UAS-CD4-tdtomato, UAS-Brp^short^-GFP, *<u>atg7</u>*: GMRflp; FRT42D, *atg7*^d4^/FRT42, GMR-Gal80; GMR-Gal4, UAS-CD4-tdtomato, UAS-Brp^short^-GFP, *<u>atg6</u>*: GMRflp; GMR-Gal4, UAS-CD4-tdtomato, UAS-Brp^short^-GFP; FRT82B, *atg6*^1^/FRT82B, tub-Gal80.

#### Immunohistochemistry and fixed imaging

Pupal and adult eye-brain complexes were dissected in cold Schneider’s Drosophila medium and fixed in 4% paraformaldehyde (PFA) in PBS for 40 minutes. Tissues were washed in PBST (0.4% Triton-X) and mounted in Vectashield (Vector Laboratories, CA). Images were obtained with a Leica TCS SP8-X white laser confocal microscope with a 63X glycerol objective (NA=1.3). The primary antibodies used in this study with given dilutions were as follows: mouse monoclonal anti-Chaoptin (1:200; Developmental Studies Hybridoma Bank); rat monoclonal anti-nCadherin (1:100; Developmental Studies Hybridoma Bank); rabbit monoclonal anti-Atg8 (1:100; Abcam); goat polyclonal anti-GFP (1:1000; Abcam); rat monoclonal anti-GFP (1:500; BioLegend); rabbit polyclonal anti-CD4 (1:600; Atlas Antibodies); rabbit polyclonal anti-DsRed (1:500; ClonTech); rabbit anti-Syd1 (1:500; gift from Sigrist Lab). The secondary antibodies Cy3, Cy5 (Jackson ImmunoResearch Laboratories) and Alexa488 (Invitrogen) were used in 1:500 dilution.

#### Brain culture and live imaging

For all ex vivo live imaging experiments an imaging window cut open removing posterior head cuticle partially. The resultant eye-brain complexes were mounted in 0.4% dialyzed low-melting agarose in a modified culture medium as described before ^13^. Live imaging was performed using a Leica SP8 MP microscope with a 40X IRAPO water objective (NA=1.1) with a Chameleon Ti:Sapphire laser and Optical Parametric Oscillator (Coherent). For single channel CD4-tdGFP imaging the excitation laser was set to 900 nm and for two-color GFP/tomato imaging lasers were set to 890 nm (pump) and 1090 nm (OPO).

#### Trans-tango and activity-dependent GRASP

For both trans-tango and GRASP experiments mosaic control and autophagy-deficient R7 photoreceptors were generated by MARCM using the combination of GMRflp and R7-specific driver Rh4-Gal4 (see “Drosophila genotypes” section for detailed genotypes). Trans-tango flies were raised at 25°C and transferred to 18°C on the day of eclosion ^36^. After 1 week of incubation at 18°C, brains were dissected and stained using a standard antibody staining protocol to label postsynaptic neurons of R7 photoreceptors. The number of postsynaptic neurons was counted manually from their cell bodies using cell counter plugin in Fiji including all cell bodies with weak or strong labelling to reveal all potential connections. For activity-dependent GRASP experiments, flies were transferred to UV-transparent Plexiglas vials on the day of eclosion and kept in a custom-made light box with UV light (25°C, 20-4 light-dark cycle) for 3 days to activate UV-sensitive R7 photoreceptors. Brains were dissected and stained with a polyclonal anti-GFP antibody to label R7 photoreceptors, monoclonal anti-GFP antibody to label GRASP signal, and polyclonal anti-CD4 antibody to label postsynaptic neurons ^43^.

#### Electroretinogram (ERG) recordings

Newly-hatched (0-day old) adult flies were collected and glued on slides using nontoxic school glue. Flies were exposed to alternating 1s “on” 2s “off” light stimulus provided by computer-controlled white LED system (MC1500; Schott). ERGs were recorded using Clampex (Axon Instruments) and quantified using Clampfit (Axon Instruments).

#### Buridan’s paradigm object orientation assay

Fly object orientation behavior was tested according to standard protocols using flies grown in low densities in a 12/12-hour light dark cycle ^32,51^. The behavioral arena consisted of a round platform of 117 mm in diameter, surrounded by a water-filled moat and placed inside a uniformly illuminated white cylinder. The setup was illuminated with four circular fluorescent tubes (Osram, L 40w, 640C circular cool white) powered by an Osram Quicktronic QT-M 1×26–42. The four fluorescent tubes were located outside of a cylindrical diffuser (DeBanier, Belgium, 2090051, Kalk transparent, 180g, white) positioned 147.5 mm from the arena center. The temperature on the platform during the experiment was 25°C and 30 mm wide stripes of black cardboard were placed on the inside of the diffuser. The retinal size of the stripes depended on the position of the fly on the platform and ranged from 8.4° to 19.6° in width (11.7° in the center of the platform). Fly tracks were analyzed using CeTrAn ^32^ and custom written python code ^51^. To reduce the complexity of the behavioral data, only absolute stripe deviation while moving was chosen, because this parameter gives a very good estimate of how precise the animals follow an object orientated path. It is calculated as an average of all points of the fly path away from an imaginary line through the two black vertical bars. For the absolute stripe deviation, it is irrelevant whether the fly deviates to the right or left. The data was statistically analyzed using ANOVA and Tukey HSD as a posthoc test using R.

### QUANTIFICATION AND STATISTICAL ANALYSIS

#### Synapse number analysis

All imaging data were analyzed and presented with Imaris (Bitplane). For synapse number analysis, CD4-tomato channel was used to generate Surfaces for individual axon terminals and Brp-positive puncta inside the Surface are filtered using the masking function. Brp-positive puncta in photoreceptor terminals were automatically detected with the spot detection module (spot diameter was set to 0.3 µ) using identical parameters between experimental conditions and corresponding controls. Synapse numbers were taken and recorded directly from statistics tab of Spot function. Graph generation and statistical analyses were done using GraphPad Prism 8.

#### Filopodia, Bulbous tip filopodia, synapse distribution analysis

All imaging data were analyzed and presented with Imaris (Bitplane). For synapse distribution analysis, Brp-positive puncta were detected following the same steps in “Synapse number analysis” in R7 axon terminals. Start and endpoints of axon terminals were selected manually with the measurements point module using nCad staining as a reference (start point=beginning of nCad staining at the most distal part of medulla (M0), end point=the beginning of M7, serpentine layer in the medulla. Note that M7 layer is devoid of synapses, hence is not labelled by nCad. The length of axon terminals are measured with the measurement point module and normalized as start point = 0 and end point = 100. The actual positions of Brp-positive puncta were exported and relative positions were calculated according to the normalized length of axon terminals. The following equation is used to calculate relative positions of Brp-positive puncta: relative position = (actual position-start point)/length x 100. For all filopodia and bulbous tip filopodia distribution analysis, the same steps were followed except that spots were manually placed on the emerging points of all visible filopodia. Graph generation and statistical analyses were done using GraphPad Prism 8.

#### Filopodia tracing

Filopodia tracing was performed as previously described ^14^. Briefly, we previously developed an extension to the Amira Filament Editor ^52^, in which an individual growth cone is visualized as annotated skeleton tree where each branch corresponds to a filopodium. In the first time step of 4D data set, the user marks the GC center, which is automatically detected in the subsequent time steps. Filopodia tips marked by the user are automatically traced from the tip to the GC center based on an intensity-weighted Dijkstra shortest path algorithm ^53^. The user visually verifies the tracing and corrects it using tools provided by the Filament Editor if necessary. After tracing all filopodia in the first time step, they are automatically propagated to the next time step with particular filopodia IDs. In every subsequent steps, the user verifies the generated tracings and adds newly emerged filopodia. This process continues until all time steps have been processed. Statistical quantities are directly extracted from the Filament Editor as spreadsheets for further data analysis.

#### Mathematical modeling

We adopted the data-driven stochastic model from ^14^. In short, the model structure remained identical, while we estimated genotype-specific parameters from the live imaging data presented in this manuscript (Figure 5b-e; Supplementary Movie 3; Table S2). In brief, we modelled synapses (S), short-lived transient bulbous tips (sB) that appeared and disappeared within the 60 minutes imaging interval and stable synaptogenic bulbous tips (synB) that persisted for more than 40 minutes. We also modelled two types of filopodia, which are distinguished by their lifetime and were denoted short-lived-(sF) and long-lived (F) filopodia.

The model’s reaction stoichiometries are determined by the following reaction scheme:

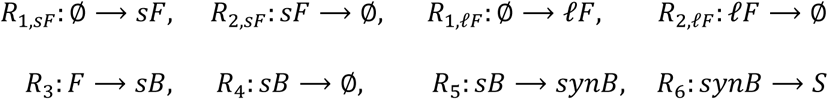

where reactions *R* _1,*sF*_ and *R* _1,*ℓF*_ denote the generation of short- and long-lived filopodia, while *R*_2, *sF*_ and *R*_2, *ℓF*_ denote their retraction. Reaction *R_3_* denotes the formation of a (transient) bulbous tip, while *R*_4_ denotes its retraction. Reaction *R*_5_ denotes the stabilization of a transient bulbous tip, and finally a stable bulb forms a synapse with reaction *R*_6_.

Note that in R_3_ we denote by F any filopodium (short-lived and long-lived) and in R_4_ we have ignored the flux back into the filopodia compartment *sF + ℓF* as it insignificantly affects the number number of filopodia (small number of bulbous tips, small rate r_4_).

Similar to the published model ^14^, reaction rates/propensities of the stochastic model are given by

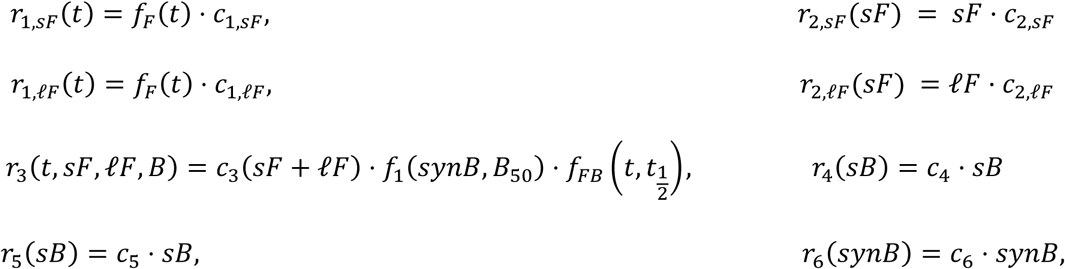

where *c*_1_ … *c*_6_ are reaction constants (estimated as outlined below). The feedback function *f*_1_(*synB, B*_50_) = (*synB* + *B*_50_)/*B*_50_ models bulbous auto-inhibition due to limited resources and synaptic seeding factor competition as introduced before ^14^. The functions *f_F_*(*t*) and 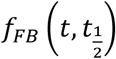 model slow-scale dynamics of filopodia- and bulbous dynamics, with previously determined parameters ^14^:

*f_FB_*(*t*) is a tanh function with

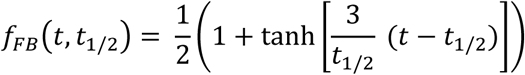

that models a time-dependent increase in the propensity to form bulbous tips with t_1/2_ = 1000 (min). The time-dependent function 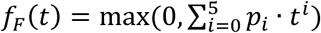 is a fifth-order polynome with coefficients p_5_ = −2.97 · 10^−14^, p_4_ = 3.31 · 10^−13^, p_3_ = −1.29 · 10^−9^, p_2_ = 2.06 · 10^−6^, p_1_ = −1.45 · 10^−3^ and p_0_ = 1 that down-regulates the generation of new filopodia at a slow time scale. Note, that *t* denotes the time in (min) after P40 (e.g. t_P40_ = 0 and t_P60_ = 60*20).

##### Parameter estimation

Using the methods explained below, we derived the parameters depicted in Table S2. We first estimated *c_2,sF_*, *c_2,ℓF_* from the filopodial lifetime data, whereby *c_2,sF_* was approximated as the inverse of the lifetimes of all filopodia that lived *less than* 8 minutes and *c_2,ℓF_* from all filopodia living *at least* 8 minutes. We realized that the number of filopodia per time instance was Poisson distributed (Supplementary Fig. 4, solid black lines), i.e. *sF∼P(A_sF_)* and *ℓF∼P(A_ℓF_)*, where λ denotes the average number of filopodia per time instance. Given the first-order retraction of filopodia (*≈* exponential lifetime), the Poisson distribution can be explained by a zero-order input with rate *c*_1,*sF*_ and *c*_1,*ℓF*_ and *A_sF_* = *r*_1,*sF*_/*c*_2,*sF*_ and *A_ℓF_ = r_1,ℓF_/c_2,ℓF_* respectively. Using the mean number of *sF*, *ℓF* at P60 we then estimated *c_1,sF_ = A_sF_(P60) · c_2,sF_/f_F_(P60)* and *c_1,ℓF_ = A_sF_(P60) · c_2,ℓF_/f_F_(P60)*.

Next, we investigated the lifetimes of bulbous tip filopodia (Supplementary Fig. 6b-e). We realized that akin to the *wild type*, the *atg6* and *atg7* exhibited almost no transient bulbous tips. We therefore set *c_4_ = 1/120* (min^−1^) according to the published model ^54^. Furthermore, we determined *c_6_* from the steepest slope in Fig. 6c (control data) divided by the average number of Bulb 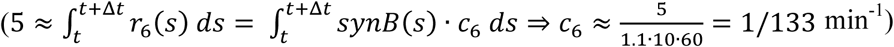. We then estimated the three parameters c_5_, B_50_ and r_3_(t) for t = P60. To do so, we used the number distribution of short-lived and synaptogenic bulbous tips (Figure 5f-g) and set up the generator matrix

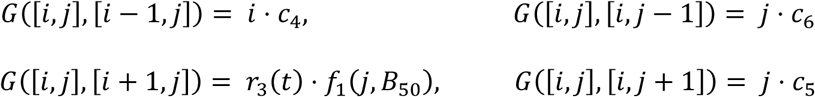

with diagonal elements such that the row sum equals 0. In the notation above, the tupel [i, j] denotes the state where *i* short-lived bulbous tips *sB* and *j* synaptogenic bulbous tips *synB* are present. The generator above has a reflecting boundary at sufficiently large N (maximum number of bulbous tips). Above, *r_3_(t)* is auto-inhibited by the number of stable bulbous tips through function *f*_1_ The stationary distribution of this model is derived by solving the eigenvalue problem

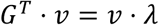

and finding the eigenvector corresponding to eigenvalue λ_0_ = 0. From this stationary distribution, we compute the marginal densities of *sB* and *synB* (e.g. summing over all states where i = 0, 1, … for sB) and fit them to the experimentally derived frequencies by minimizing the Kullback-Leibler divergence between the experimental and model-predicted distributions. Lastly, parameter is derived by calculating

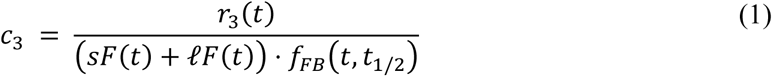

where *sF(t) = sF(t_P_*_60_), *ℓF(t) = ℓF(t_P_*_60_) and *f_FB_(t) = f_FB_(t_P_*_60_, *t*_1/2_).

##### Mechanistic model explains autophagy mutant phenotypes as a consequence of increased seeding factor abundance

We adopted the mechanistic model from ^14^. This model essentially assumes a dynamic pool of a limited resource of bulbous-tip stabilizing factors (Fig. 6g; Supplementary Fig. 6a). The model consists of four types of reactions: new filopodia emerge (reaction *G_1_*), accumulate resources (reaction *G_2_*), retract (reaction *G_3_*) or release resources (reaction *G_4_*).

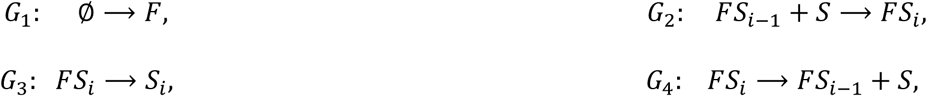

where *F* denotes an ‘empty’ filopodium, *S* denotes the seeding factor and *FS*_i_ denotes a filopodium with *i* seeding factor proteins in it. The reaction rates (propensities) were modelled as

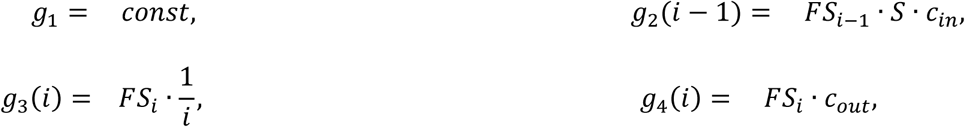

where we set *g*_1_ equal to the average rate of transient bulbous tip emergence in the control experiments at P60, i.e. *g_1_ = r_3_(t_P60_, WT)*. Reaction rate *g*_3_ implements a competitive advantage: the lifetime of bulbous filopodia is increased proportionally to the number of seeding factors it accumulated. The parameters *c_in_* and *c_out_* were set to values 0.07 and 1.5 (time^−1^) and as initial condition we set 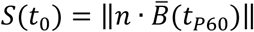, where *n* is the number of states (we used *n* = 120), 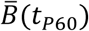 denotes the genotype-specific average number of bulbous tips at P60 and ‖·‖ denotes the next integer function.

Importantly, in the model, the wildtype and the *atg6*- and *atg7*-knockout mutants only differ in the total number of seeding factors available.

We stochastically ran the model 100,000 timesteps to reach a steady state and discarded the first half as a burn-in period (pre-steady state). Subsequently, we analyzed the number of bulbous tips and their lifetimes from the remaining time steps as shown in Supplementary Fig. 6b-i. Thereby, we assumed that filopodia would be recognized as bulbous tips only if they contained at least n/4 seeding factors.

In summary, these computational experiments highlight that the phenotype of the *atg6*- and *atg7*-knockout mutants can be solely explained by an increased abundance of seeding factors (= compromised ability to degrade seeding factors).

In the case of autophagy upregulation *(atg6*, GMR>Atg6), we observed a different phenotype: From the data-driven model we could see that bulbous tips were destabilized (parameter r4 in Supplementary Table 2), and also that the feedback was lost (parameter E[f1] close to 1 in Supplementary Table 2). We tested different parameter- and model alterations to reproduce both the number- and life time distribution of bulbous tips. Finally, we found that if seeding factors no longer stabilized bulbous tips (loss in the competitive advantage), both the life time- and the number distribution of bulbous tips can be accurately reproduced. Thus, we set reaction rate *g*_3_ to *g_3_ = FS · canst*, for autophagy upregulation, where *canst = c_4_*(time^−1^; Supplementary Table 3).

### DATA AND CODE AVAILABILITY

Raw (.lif format) and processed (.ims and .am format) imaging datasets are available on request. The filopodia tracking software is an extension of the commercial software Amira, which is available from Thermo Fisher Scientific. The filopodia tracking software is available from the corresponding author upon request in source code and binary form. Executing the binary requires a commercial license for Amira. MATLAB codes for model parameter inference for model simulation have previously been published ^14^ and are available through https://github.com/vkleist/Filo.

### LEAD CONTACT AND MATERIALS AVAILABILITY

All reagents used in this study are available for distribution. Requests for resources and reagents should be directed to Robin Hiesinger.

## Supplemental Information

**Supplementary Figure 1.**
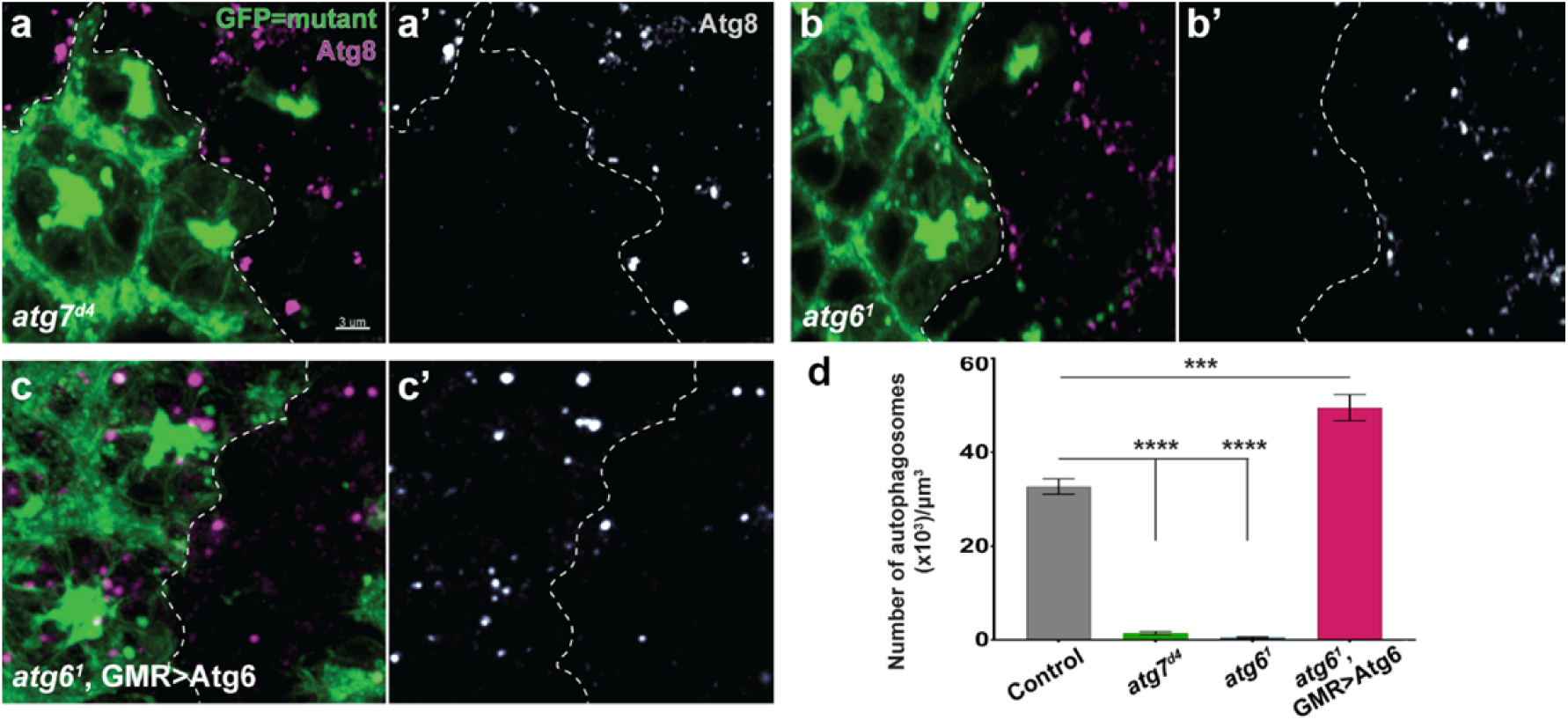
Atg6 and Atg7 are required for developmental autophagy in *Drosophila* photoreceptors. **a-c’,** Atg8 immunolabelled autophagosomes in GFP-positive photoreceptor clones of *atg7*^d4^ **(a-a’)**, *atg6*^1^ **(b-b’)**, and *atg6*^1^, GMR>Atg6 **(c-c’)** versus non-GFP control clones in genetic mosaics of P+50% pupal retina. **d,** Number of autophagosomes in a given volume. Note almost complete abolishment of autophagosomes in *atg7*^d4^ and *atg6*^1^ mutant photoreceptors and a significant increase in autophagosome number in *atg6*^1^, GMR>Atg6 photoreceptors. n=8 retinas per condition, one region of interest is randomly selected per retina. Unpaired t-test; ***p<0.001, ****p<0.0001. Error bars denote mean ± SEM.

**Supplementary Figure 2.**
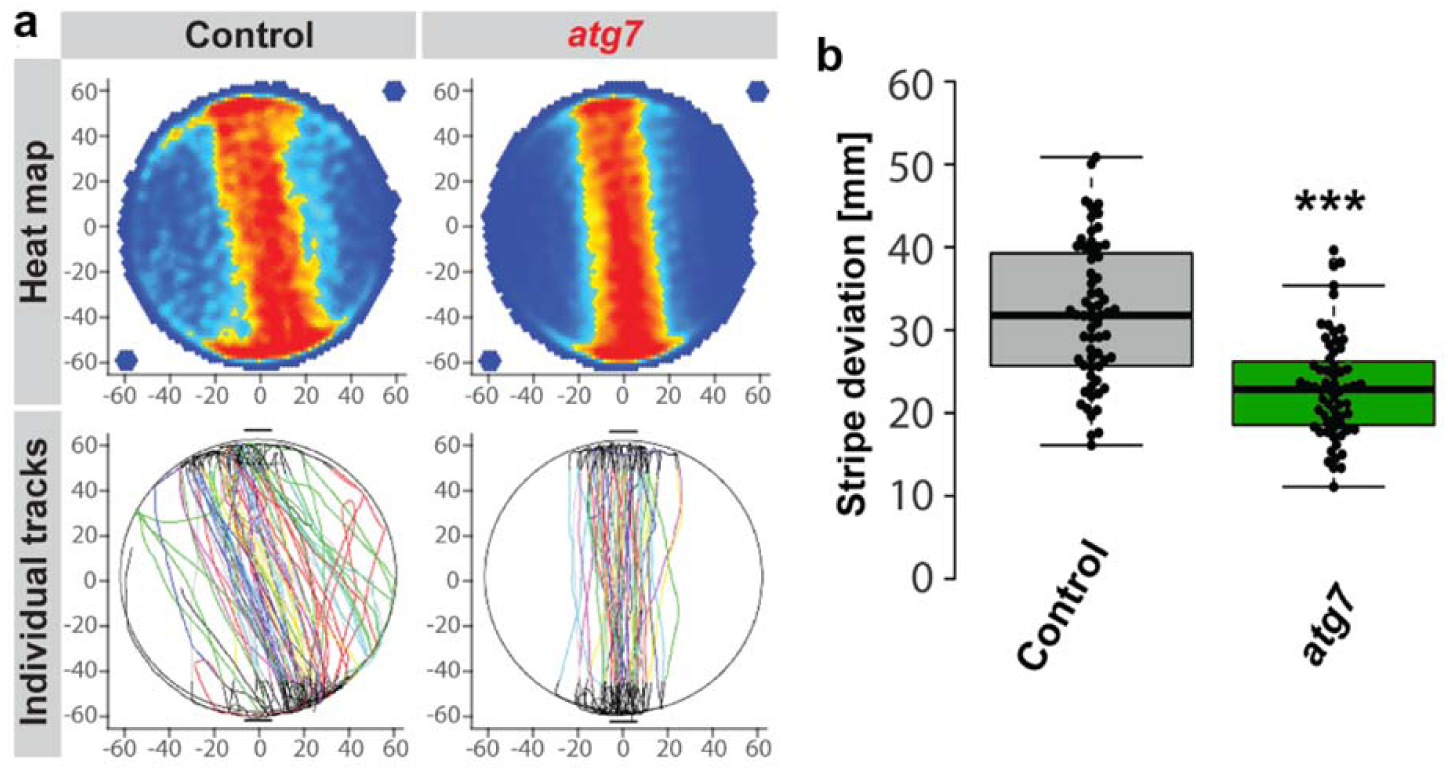
Loss of *atg7* in *Drosophila* photoreceptors leads to increased visual attention behavior. **a,** Stripe fixation behavior of adult flies with control and *atg7* mutant photoreceptors is shown on the population level (heatmap) and as individual tracks. **b,** Quantification of stripe deviation. n=60 flies per condition, two-way ANOVA and Tukey HSD as post hoc test, ***p<0.001. Note that similar to flies with *atg6* mutant photoreceptors (see Fig. 1h), flies with *atg7* mutant photoreceptors show increased stripe fixation behavior and repetitive walks between stripes.

**Supplementary Figure 3.**
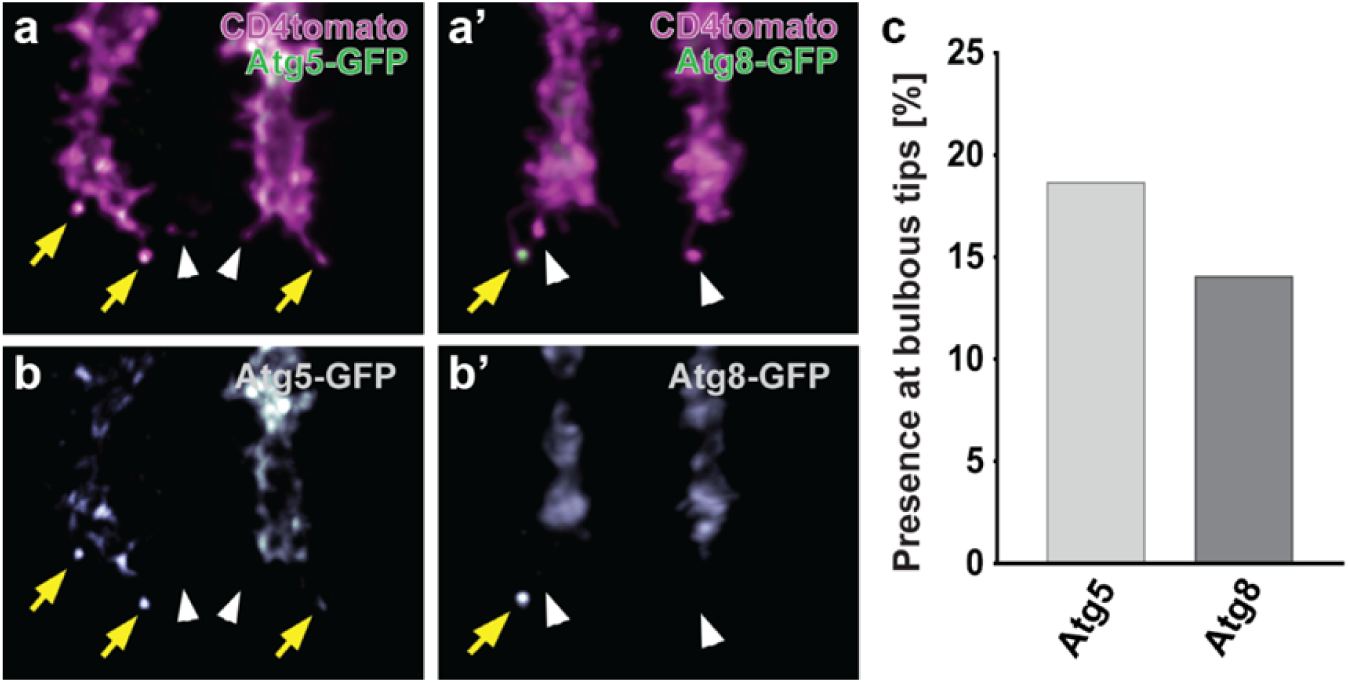
The essential autophagy proteins Atg5 and Atg8 localize to synaptogenic filopodia tips. **a-b’,** Localization of autophagy essential proteins Atg5 (**a-a’**) and Atg8 (**b**-**b’**) to bulbous tip filopodia (P+60%). Yellow arrows show the presence of Atg5 and Atg8 at bulbous tips, while white arrowheads show bulbous tips without Atg5 and Atg8. **c,** Percentage of bulbous tip filopodia with Atg5 and Atg8 signal to all bulbous tip filopodia. n=30 terminals. All bulbous tip filopodia from 30 axon terminals were pooled for quantification.

**Supplementary Figure 4.**
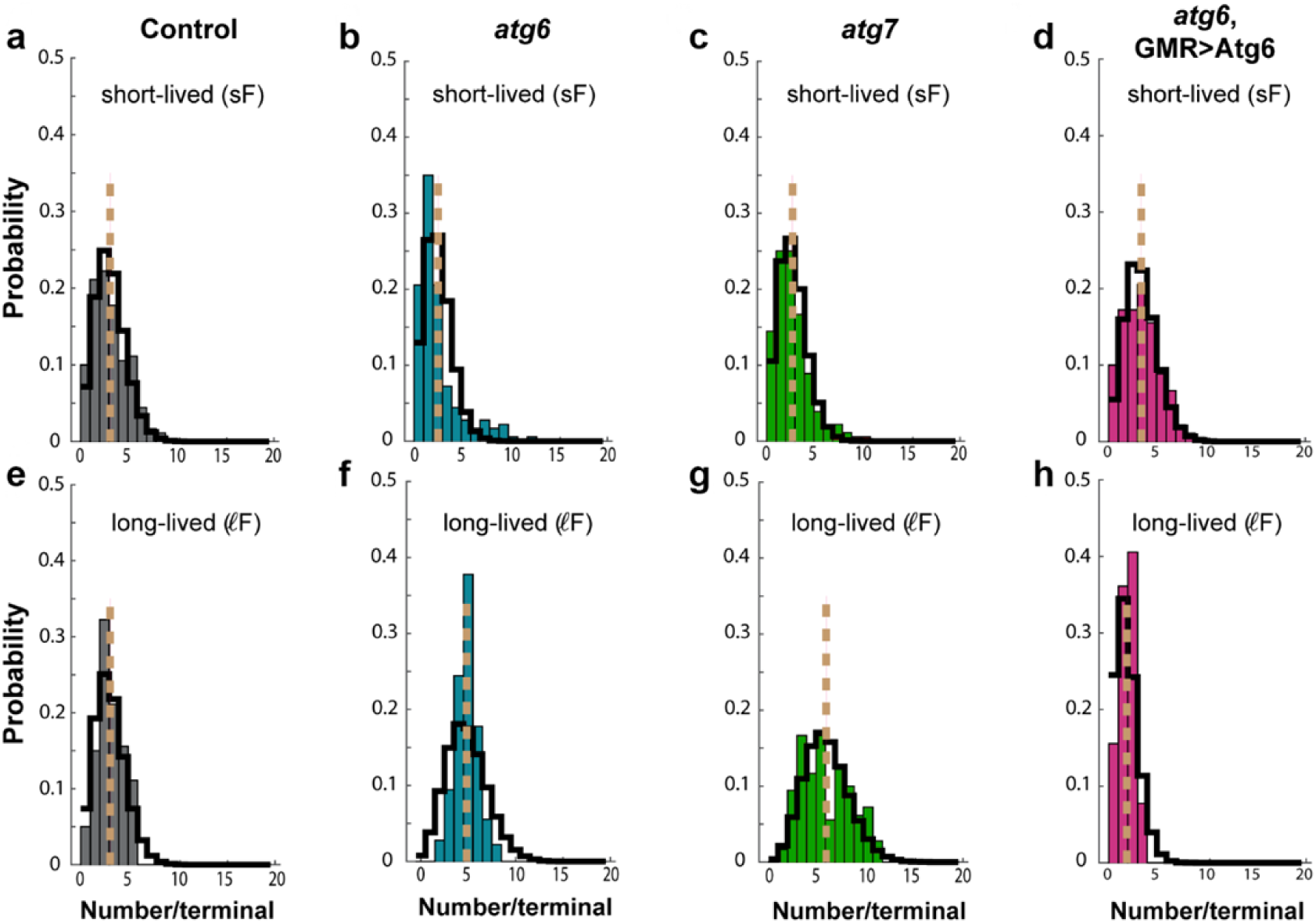
Number of short-lived and long-lived filopodia at P60. Bars denote the observed numbers during live imaging and the dashed vertical line indicates the average numbers. The solid black trace depicts a Poisson distribution with expectation value equal to the average number of observed filopodia. short-lived filopodia = filopodia exist shorter than 8 mins, long-lived filopodia = filopodia exist longer than 8 mins. Values for lifetimes and numbers are shown in Supplementary Table 1.

**Supplementary Figure 5.**
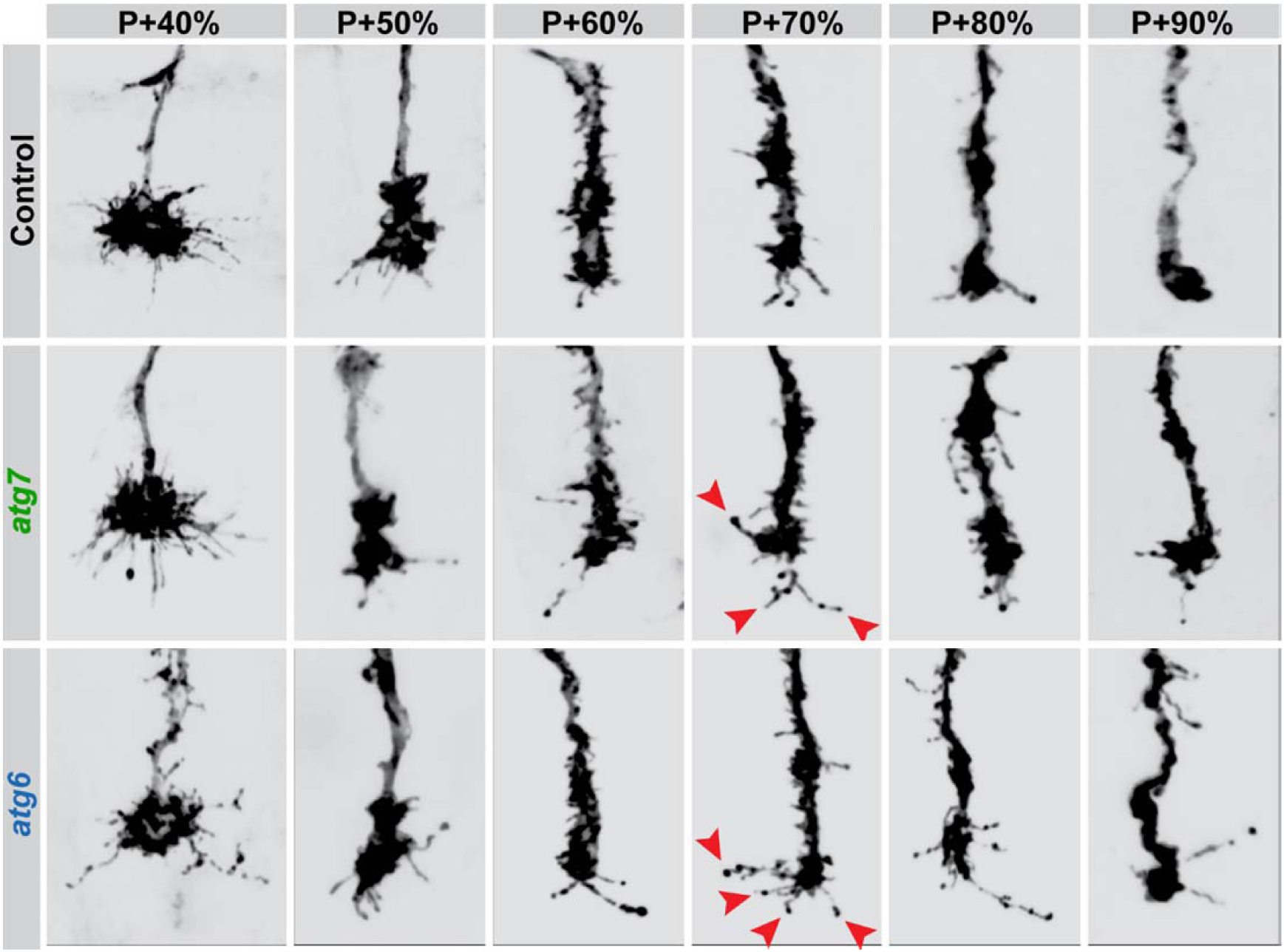
Morphology of R7 photoreceptor axon terminals throughout the second half of pupation (the period of synapse formation). Representative images of control, atg7, and atg6 mutant R7 axon terminal morphologies at P+40%, P+50%, P+60%, P+70%, P+80%, and P+90% pupal development. Red arrowheads show examples of supernumerary bulbous tip filopodia at P+70%. Note that loss of autophagy leads to increased numbers of bulbous tip filopodia especially during the peak time of synaptogenesis (P+60%-P+80%).

**Supplementary Figure 6.**
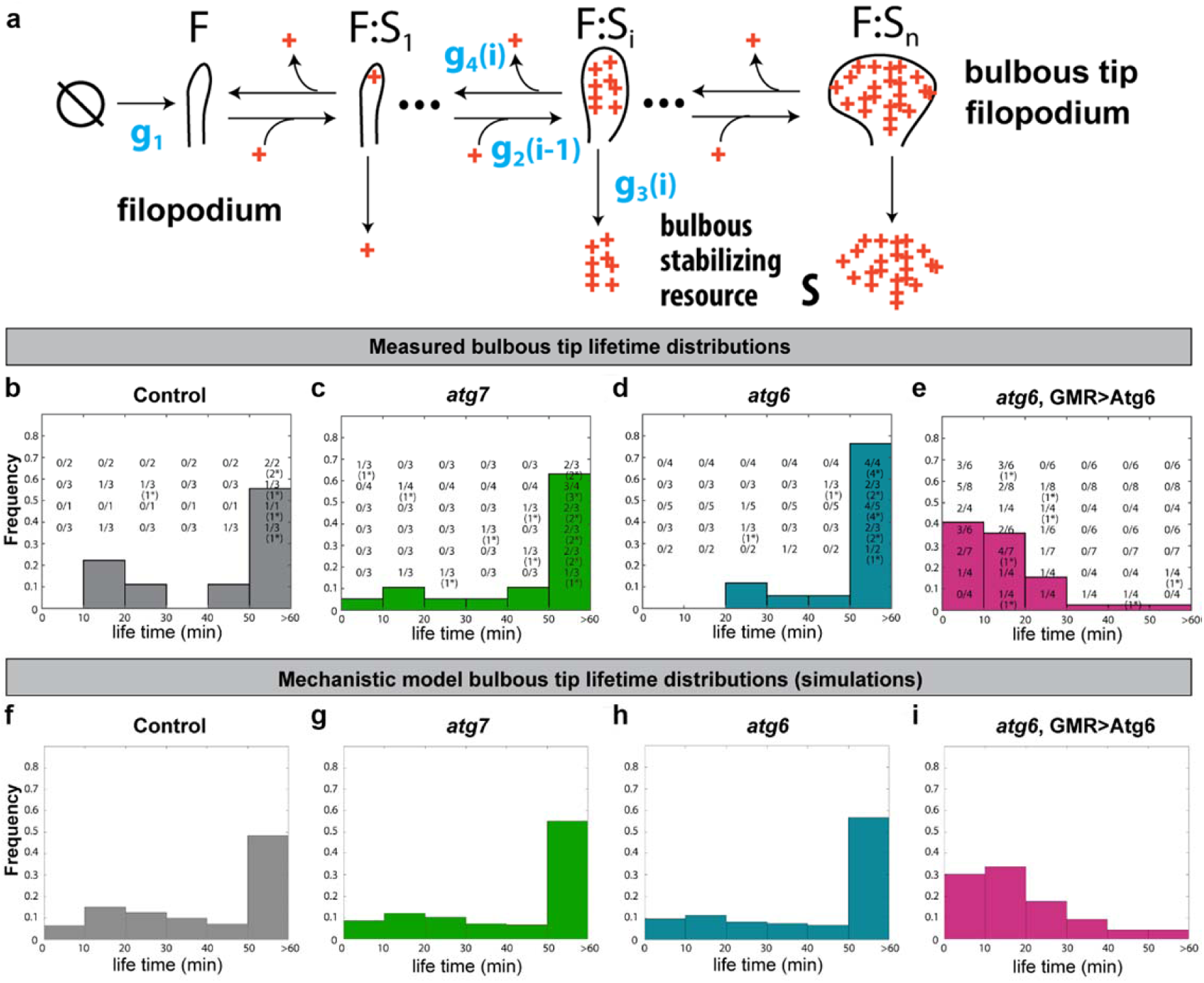
The Mechanistic Model: Lifetimes of synaptogenic bulbous tip filopodia as a function of a limiting resource of synaptic seeding factors. **a,** Graphical depiction of the mechanistic model. **b-e,** <u>Measured data:</u> Histograms depicting the observed frequency of the respective bulbous tip life times during live imaging at P60. The numbers on histograms indicate the number of observations in the respective life time category per growth cone. Numbers in brackets with a star, e.g. (1*), indicate that the bulbous tip either already existed in the first imaging frame, or persisted until the last image. Thus, these life times might actually be longer than indicated here. **f-i,** <u>Model output:</u> Histograms depicting the frequency of the respective bulbous tip lifetimes according to simulations using the mechanistic model. Note that the mechanistic model successfully recapitulates the observed lifetimes of bulbous tip filopodia.

**Supplementary Figure 7.**
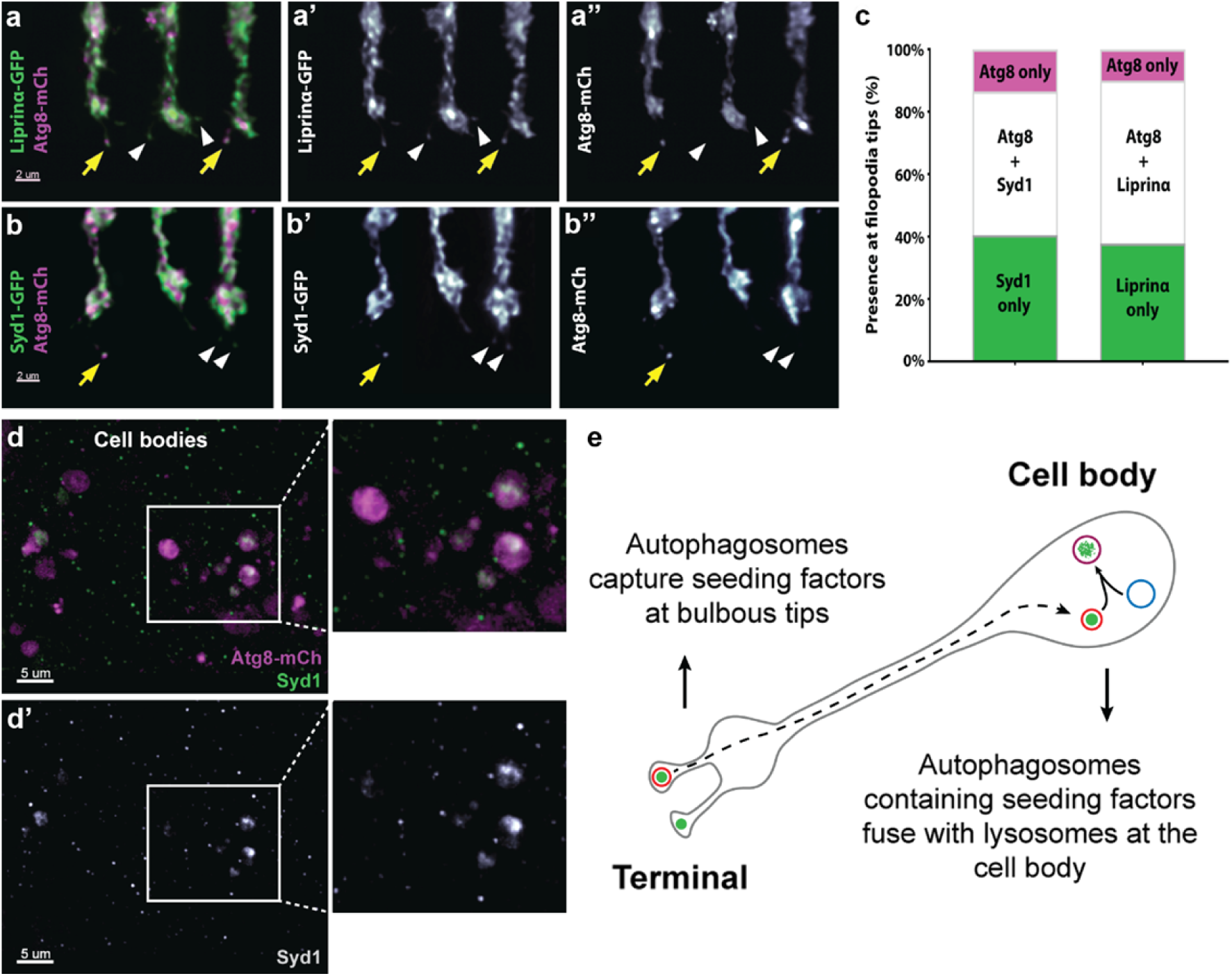
Autophagosomes colocalize with synaptic seeding factors at filopodia tips and contain synaptic seeding factors in large degradative multivesicular compartments at cell bodies. **a-a’’,** Representative R7 axon terminals expressing Liprin-α-GFP and Atg8-mCherry. **b-b’’,** Representative R7 axon terminals expressing Syd-1-GFP and Atg8-mCh. Yellow arrows: co-localization of Atg8 with synaptic seeding factors Liprin-α and Syd-1 at filopodia tips; white arrowheads: Liprin-α and Syd-1 at filopodia tips without apparent Atg8 co-localization. **c,** Percentages of Syd-1 only, Liprin-α only, Atg8 and Syd-1 together (Atg8 + Syd-1), Atg8 and Liprin-α together (Atg8 + Liprin-α), and Atg8-only filopodia tips. n=30 terminals per condition. Note that most Atg8-positive compartments are also positive for the synaptic seeding factors. All filopodia from 30 terminals were pooled for quantification. **d-d’,** Atg8-positive multivesicular vacuoles contain endogenous Syd1 (detected with anti-Syd1 antibody) at photoreceptor cell bodies. **e,** Schematic of proposed mechanism of degradation of synaptic seeding factors by autophagy in photoreceptor neurons, including capture at axon terminal filopodia tips and degradation during retrograde transport to the cell body, as first shown in vertebrate cell culture ^44^.

**Table S1:** Lifetimes (min) and average numbers of short- and long-lived filopodia at P60. Mean ± (standard deviation). Number distributions are shown in Supplementary Fig. 4.

| | Short lived (sF) | | Long-lived ($\ell F$) | |
| --- | --- | --- | --- | --- |
|  | Life time | Number | Life time | Number |
| wild type | 2.3 (1.6) | 2.6 (1.8) | 15 (10) | 2.6 (1.4) |
| *atg6* | 2.7 (1.9) | 2 (2.3) | 21 (16) | 4.8 (1.2) |
| *atg7* | 2.4 (1.6) | 2.2 (1.9) | 20 (15) | 5.6 (2.6) |
| *atg6*, GMR>Atg6 | 2.2 (1.7) | 2.9 (1.9) | 13 (7) | 1.4 (0.84) |

**Table S2:** Measured average rates of the data-driven model at P60. The denotation is taken from the original model in Figure 3A of ^14^ and refer to the following filopodial transitions:

|  | r3 | r2B | E[f1] | r4 | r5 | Avg. bulbs |
| --- | --- | --- | --- | --- | --- | --- |
| wild type | 0.0122 | 0.0948 | 0.1291 | 0.0014 | 0.0108 | 1.653 |
| *atg6* | 0.0229 | 0.0932 | 0.2463 | 0.0025 | 0.0205 | 3.028 |
| *atg7* | 0.0189 | 0.1985 | 0.0955 | 0.0019 | 0.0170 | 2.501 |
| *atg6*, GMR>Atg6 | 0.1032 | 0.1085 | 0.9515 | 0.1010 | 0.0018 | 1.644 |

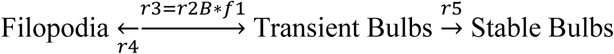 r3: measured rate of bulb formation, contains r2B * f1, unit: 1/min r2B: propensity to form bulbs, cannot be measured, because feedback f1 reduces r2B, shown is the only possible fit of r2B, unit: 1/min f1: negative feedback on bulb formation, cannot be measure, see r5, shown is the only possible fit of the data (r2B; smaller f1 indicates stronger feedback; f1=1 indicates no feedback r4: measured rate of bulb disappearance, unit: 1/min r5: measured rate of bulb stabilization, unit: 1/min Avg. bulbs: average number of bulbs per time instance (min) over an hour (P60) In blue: direct measurements

**Table S3:** Parameters of the data-driven model. All parameters in units min^−1^ except for B_50_ (unitless) and t_1/2_ (min). ^§^previously determined ^14^.

| | $c_{1,sF}$ | $c_{2,sF}$ | $c_{1,\ell F}$ | $c_{2,\ell F}$ | $c_3$ | $c_4$ | $c_5$ | $c_6$ | $B_{50}$ | $t_{1/2}$ |
| --- | --- | --- | --- | --- | --- | --- | --- | --- | --- | --- |
| wild type | 1.82 | 0.43 | 0.28 | 0.07 | 0.024 | 1/120<sup>§</sup> | 0.063 | 1/133 | 0.078 | 1000<sup>§</sup> |
| *atg6* | 1.19 | 0.37 | 0.37 | 0.05 | 0.018 | 1/120<sup>§</sup> | 0.068 | 1/133 | 0.716 | 1000<sup>§</sup> |
| *atg7* | 1.48 | 0.42 | 0.45 | 0.05 | 0.033 | 1/120<sup>§</sup> | 0.075 | 1/133 | 0.162 | 1000<sup>§</sup> |
| *atg6*, GMR>Atg6 | 2.13 | 0.45 | 0.17 | 0.08 | 0.033 | 0.071 | 0.001 | 1/133 | 3.733 | 1000<sup>§</sup> |

